# The mechanism of *Gα_q_* regulation of *PLCβ3*-catalyzed *PIP2* hydrolysis

**DOI:** 10.1101/2023.08.29.555394

**Authors:** Maria E. Falzone, Roderick MacKinnon

**Affiliations:** Laboratory of Molecular Neurobiology and Biophysics, The Rockefeller University, New York, United States; Howard Hughes Medical Institute, The Rockefeller University, New York, United States

**Author notes:** **Author Contributions:** R.M. and M.E.F. designed studies; M.E.F. performed experiments; R.M. and M.E.F. analyzed and interpreted data; R.M. and M.E.F. wrote the paper. **Competing Interest Statement:** The authors declare no competing interests.

**Keywords:** *PLCβ3*, *Gα_q_*, *PIP2*, *Gβγ*, GPCR signaling

## Abstract

*PLCβ* enzymes cleave *PIP2* producing IP3 and DAG. *PIP2* modulates the function of many ion channels, while IP3 and DAG regulate intracellular Ca^2+^ levels and protein phosphorylation by protein kinase C, respectively. *PLCβ* enzymes are under the control of GPCR signaling through direct interactions with G proteins *Gβγ* and *Gα*_*q*_ and have been shown to be coincidence detectors for dual stimulation of *Gα*_*q*_ and G*α*_i_ coupled receptors. *PLCβs* are aqueous-soluble cytoplasmic enzymes, but partition onto the membrane surface to access their lipid substrate, complicating their functional and structural characterization. Using newly developed methods, we recently showed that *Gβγ* activates *PLCβ3* by recruiting it to the membrane. Using these same methods, here we show that *Gα*_*q*_ increases the catalytic rate constant, *k*_*cat*_, of *PLCβ3*. Since stimulation of *PLCβ3* by *Gα*_*q*_ depends on an autoinhibitory element (the X-Y linker), we propose that *Gα*_*q*_ produces partial relief of the X-Y linker autoinhibition through an allosteric mechanism. We also determined membrane-bound structures of the *PLCβ3-Gα*_*q*,_ and *PLCβ3-Gβγ(2)-Gα*_*q*_ complexes, which show that these G proteins can bind simultaneously and independently of each other to regulate *PLCβ3* activity. The structures rationalize a finding in the enzyme assay, that co-stimulation by both G proteins follows a product rule of each independent stimulus. We conclude that baseline activity of *PLCβ3* is strongly suppressed, but the effect of G proteins, especially acting together, provides a robust stimulus upon G protein stimulation.

**Significance Statement:** For certain cellular signaling processes, the background activity of signaling enzymes must be minimal and stimulus-dependent activation robust. Nowhere is this truer than in signaling by *PLCβ3*, whose activity regulates intracellular Ca^2+^, phosphorylation by Protein Kinase C, and the activity of numerous ion channels and membrane receptors. In this study we show how *PLCβ3* enzymes are regulated by two kinds of G proteins, *Gβγ* and *Gα*_*q*_. Enzyme activity studies and structures on membranes show how these G proteins act by separate, independent mechanisms, leading to a product rule of co-stimulation when they act together. The findings explain how cells achieve robust stimulation of *PLCβ3* in the setting of very low background activity, properties essential to cell health and survival.

## Introduction

*Phospholipase Cβ* (*PLCβ*) enzymes cleave *PIP2* in the plasma membrane to produce inositol triphosphate (IP3) and diacylglycerol (DAG), (1, 2). *PIP2* regulates the function of many membrane proteins including ion channels, IP3 increases cytoplasmic Ca^2+^ via the IP3 receptor, and DAG activates protein kinase C, which itself regulates numerous target proteins (3-5). Because *PIP2*, IP3 and DAG are critical to so many cellular processes, their tight regulation by *PLCβs* is essential to normal cellular function. *PLCβ* enzymes are under the control of G protein coupled receptor (GPCR) signaling through direct interaction with G proteins, *Gα*_*q*_ and *Gβγ* (6-8). Basal activity of *PLCβs* is maintained at very low levels in cells via two autoinhibitory elements, the X-Y linker, which occupies the active site, and the H*α*2’ in the proximal c-terminal domain (CTD), whose mechanism of autoinhibition is not well-understood (9-13).

*PLCβs* are aqueous soluble enzymes that must partition onto the membrane to carry out *PIP2* hydrolysis, which has posed a challenge to obtaining a quantitative description of their catalysis and regulation by G proteins. To overcome this challenge, we recently developed methods to measure both the partitioning of *PLCβ* enzymes between aqueous solution and membrane surfaces and the hydrolysis of *PIP2* by membrane-bound enzyme (14). With these methods, we showed that *PLCβ3* is a Michaelis-Menten enzyme and that *Gβγ-*dependent activation is mediated by increasing its local concentration at the membrane surface. *Gβγ* does not significantly change the catalytic rate constant (*k*_*cat*_) of *PLCβ3*, nor does it alter its autoinhibitory elements in structures of the *PLCβ3-Gβγ* complex (14).

The mechanism of activation by *Gα*_*q*_ is not understood, particularly the potential role of the membrane in activation. Specifically, it is not clear whether *Gα*_*q*_ activates by membrane recruitment like *Gβγ* or whether it increases *k*_*cat*_ through an allosteric mechanism. The lipid anchor on *Gα*_*q*_ is not required for activation of *PLCβs*, in contrast to *Gβγ*, suggesting that membrane recruitment might not underlie *Gα*_*q*_-dependent activation (10, 11, 15, 16). However, non-lipidated *Gα*_*q*_ has been shown to maintain its association with membranes in cells and *in vitro*, raising the possibility that membrane recruitment could still play a role even in the absence of a covalent lipid group (15). It has also been established that *Gα*_*q*_ does not activate *PLCβs* in the absence of a membrane environment, suggesting that the membrane does play a role in activation (13).

Structural studies of the *PLCβ3-Gα*_*q*_ complex in the absence of membranes revealed that *Gα*_*q*_ binds to the proximal and distal CTD of *PLCβ3* and *Gα*_*q*_ binding displaces the autoinhibitory H*α*2’ away from its binding site on the catalytic core by ~50 Å (10, 16). These observations led to the proposal that *Gα*_*q*_ actives *PLCβs* by relieving H*α*2’ autoinhibition. However, the mechanism of autoinhibition by H*α*’ is unknown: it is only known that removing H*α*2’ or disrupting its contacts with the catalytic core increases the basal activity of *PLCβs* (9, 11).

Some *PLCβs* can also be activated by *Gβγ* and *Gα*_*q*_ simultaneously. This dual activation, which underlies many physiological functions, was proposed to play a role in coincidence detection under co-stimulation of G*α*_i_ and *Gα*_*q*_-coupled receptors in cells (8, 17, 18). Dual activation was proposed to be synergistic, or greater than the sum of the activation of each G protein on its own (19).

The goal of the present study is to understand the mechanism of activation of *PLCβ3* by *Gα*_*q*,_ and of dual activation by *Gα*_*q*_ and *Gβγ*. Using functional experiments, membrane partitioning studies, and structural studies on membrane surfaces, we show that soluble *Gα*_*q*_ activates *PLCβ3* by increasing its catalytic rate constant, *k*_*cat*_, without affecting membrane recruitment. We also show that *Gα*_*q*_-stimulated enhancement of *k*_*cat*_ is mediated by the X-Y linker autoinhibitory element. Thus, the X-Y linker is a suppressor of *k*_*cat*_ that is partially relieved by *Gα*_*q*._ Finally, we show that *Gα*_*q*_ and *Gβγ* regulate *PLCβ* function independently, the former through *k*_*cat*_ and the latter through membrane recruitment. Consequently, dual stimulation yields activity enhancement equal to the product of the two independent stimuli.

## Results

### Activation of PLCβ3 by non-lipidated Gα_q_

*PLCβ3* is an aqueous soluble enzyme that partitions onto the membrane surface to catalyze *PIP2* hydrolysis. As we will describe below, we use the partition coefficient of *PLCβ3* to calculate its membrane surface concentration from its aqueous concentration set by experimental design (14). For reasons that will become apparent, we use two different concentration units in this study. In some circumstances we specify molar concentration using the notation [*quantity*]_*molar*_. In other circumstances we specify mole fraction (*mf*) in *mole%* (100 x *mf*) using the notation [*quantity*]. Because moles of solvent (water for aqueous solutions and lipids for membranes) are in vast excess of moles of solute (*PLCβ3* for aqueous solutions and *PLCβ3* and *PIP2* for membranes), we approximate *mf* as moles solute over moles solvent (14).

To measure *PIP2* hydrolysis by *PLCβ3* on membrane surfaces we employed an enzyme assay described in a recent publication (14). Briefly, the reaction takes place on a planar lipid bilayer formed over a hole in a partition separating two aqueous chambers (Fig. 1A). The lipid bilayer is comprised of 2DOPE:1POPC:1POPS (wt:wt:wt), plus a pre-determined concentration of *PIP2*, ([*PIP2*] = 1.0 *mol%*) and a *PIP2*-gated ion channel is incorporated into the bilayer via vesicle fusion. The ion channel, a modified *PIP2*-dependent, G protein-dependent inward rectifier K^+^ channel, called GIRK-ALFA, is calibrated so that the normalized K^+^ current level can be converted to membrane *PIP2* concentration (Fig. 1B-D)(14). Upon addition of *PLCβ3* under continuous mixing, after an approximately 2 s delay, the GIRK-ALFA current decreases due to hydrolysis of *PIP2* by the added *PLCβ3* (Fig. 1B). Using the predetermined calibration curve (Fig. 1C), normalized current as a function of time (Fig. 1B) can be converted to *PIP2* concentration as a function of time, as shown (Fig. 1D). The latter curve corresponds very well (typical R^2^ > 0.9) to a Lambert W Function (aka Product Log Function)(20), which describes a linear decay initially (when *PIP2* concentration is high) and an exponential decay at later times (when *PIP2* concentration is low) (Fig. 1D). The Lambert W Function derives from integration of the well-known Michaelis-Menten enzyme equation,

**Figure 1:**
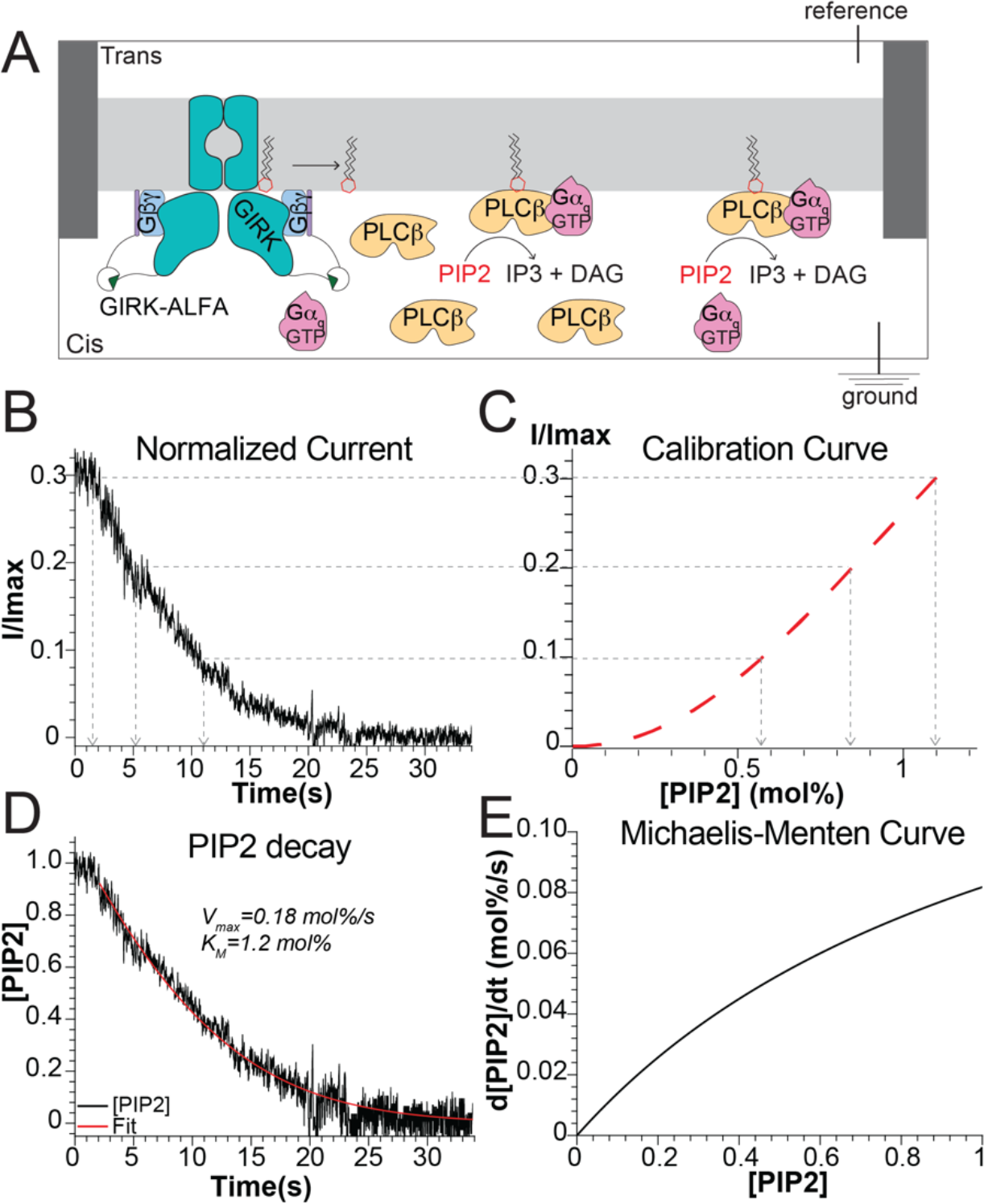
Summary of *PLCβ* functional assay and analysis. A: Cartoon schematic of planar lipid bilayer setup used to measure *PLCβ3* function. B-D: Summary of analysis of current *PLCβ3*-dependent current decays. The *PIP2* calibration curve for GIRK-ALFA (C) is used to convert the normalized current decay (B) to *PIP2* decay (D). Points on the normalized current decay are matched to [*PIP2*] and time. The resulting *PIP2* decay as a function of time is fit (R^2^=0.97) to the Lambert W Function (Eq. 8) derived through integration of the Michaelis-Menten equation with free parameters *V*_*max*_ and *K*_*M*_ shown. E: Corresponding Michaelis–Menten curve with the *K*_*M*_ and *V*_*max*_ values determined in (D).

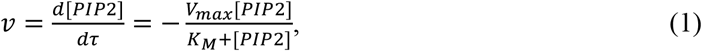

which, when integrated from *τ* = 0 to *τ* = t, yields

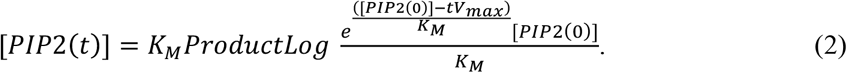

An instructive description of the relationship between the Michaelis-Menten equation (Eq. 1) and the Lambert W Function (Eq. 2) and the suitability of the latter to our studies is given in Appendix 1 of Supplementary Information. In practice, we fit the normalized current data, i.e., Fig. 2A, 3, S2, directly with a function that is the Lambert W Function transformed by the calibration curve that converts normalized current to *PIP2* concentration (Fig. 1C and Eq. 8). This function has three free parameters, *V*_*max*_, *K*_*M*_ and *C*, the latter a dimensionless (*I/I*_*max*_) current accounting for the small (almost inconsequential) nonspecific ‘leak’ current observed at the longest recorded times (Fig. 1B and 2A, S2A). This enzyme assay permits reproducible estimates of *V*_*max*_ and *K*_*M*_ for *PLCβ3* over a wide range of experimental conditions (14).

**Figure 2:**
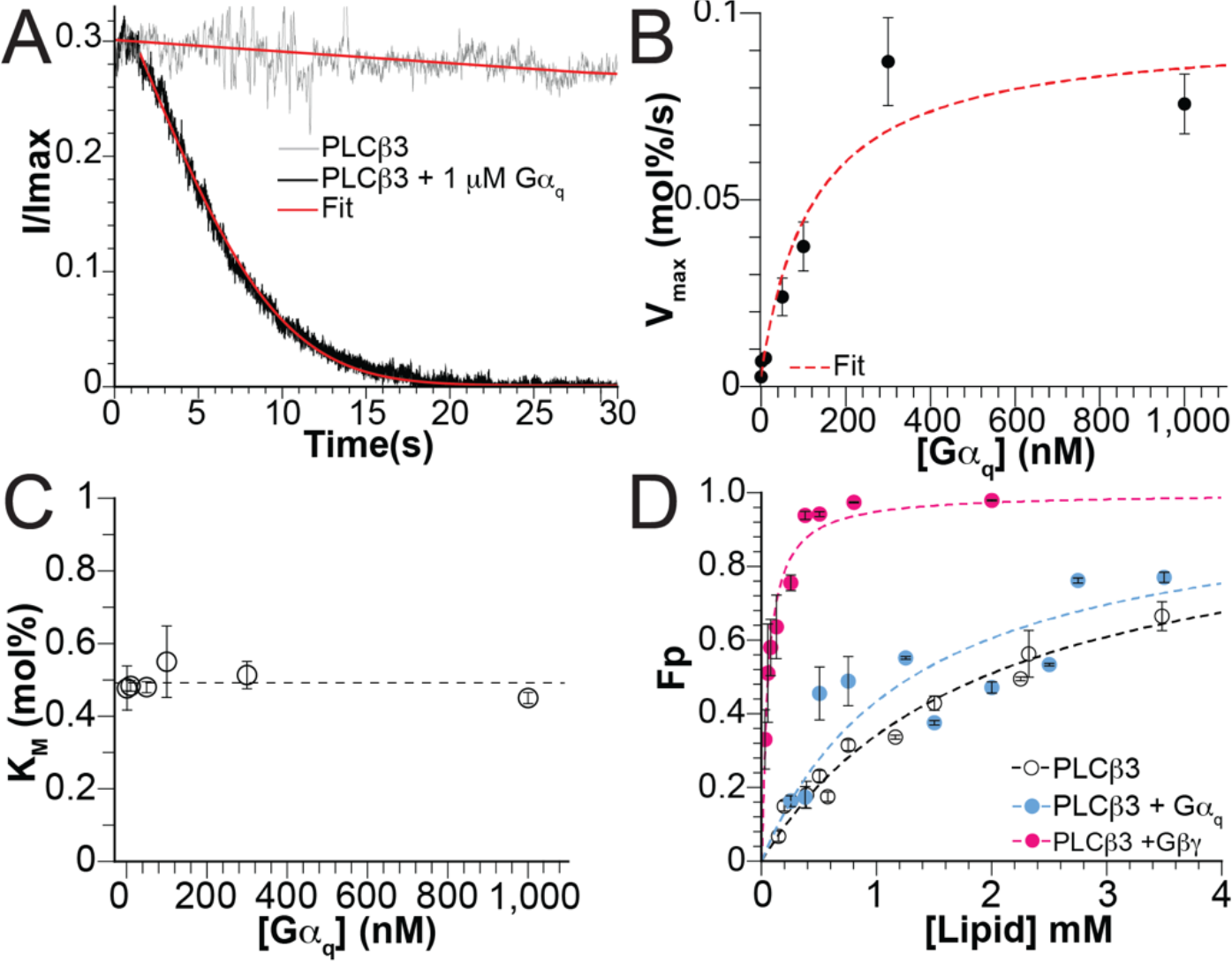
Activation of *PLCβ3* by non-lipidated *Gα*_*q*_. A: Representative *PLCβ3*-dependent normalized current decay with 29 *nM* enzyme in the absence of *Gα*_*q*_ (gray) or in the presence of 1.0 *µM Gα*_*q*_ (black) fit to Eq. 8 (red curve). In the presence of *Gα*_*q*,_ *V*_*max*_=0.091 *mol%/s, K*_*M*_=0.42 *mol%*, C= 0.00092, R^2^=0.99. B: *V*_*max*_ as a function of *Gα*_*q*_ concentration fit to 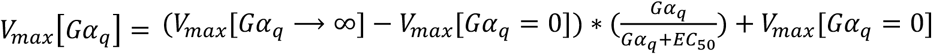 and EC_50_ where *V*_012_[*Gα*_A_ = 0] is the *V*_*max*_ in the absence of *Gα*_*q*_, 0.0026 *mol%/s* (14). EC_50_=120 *nM* and *V*_012_>*Gα*_A_ → ∞A =0.095, R^2^=0.91. Individual points are from at least 3 repeats and the error bars are standard error of mean. C: *K*_*M*_ as a function of *Gα*_*q*_ concentration. Dashed line highlights the mean of all values. Individual points are from at least 3 repeats and the error bars are standard error of mean. D: Membrane partitioning curve for wildtype *PLCβ3* in the presence of *Gα*_*q*_ Q209L (blue) for 2DOPE:1POPC:1POPS LUVs with Fraction Partitioned (*F*_*p*_) on the Y axis. Points are the average from 2 repeats for each lipid concentration and error bars are range of mean. Data were fit to Eq. 5 to determine *K*_*x*_ (dashed blue curve). *K*_*x*_=4.2*10^4^, R^2^=0.68. Data points and the fit to Eq. 5 for *PLCβ3* alone (black) and in the presence of *Gβγ* (pink) are shown for reference (14).

To ensure that *PLCβ3* was not affected by product inhibition under our assay conditions, we tested its function in the presence of *Gβγ* and 1.0 *mol%* DAG or 1.0 *µM* IP3 (Fig. S3). Current decays and determined values for *V*_*max*_ and *K*_*M*_ are indistinguishable from experiments without DAG and IP3 (Fig. S3), indicating that *PLCβ3* is not inhibited by the products of its catalyzed reaction in our experimental setup.

To study the activation of *PLCβ3* by *Gα*_*q*_ we used non-lipidated *Gα*_*q*_ (10, 11, 15). Because the GTP bound form of *Gα*_*q*_ is required to activate *PLCβ3*, we used a hydrolysis-deficient mutant (Q209L) that remains constitutively bound to GTP (Fig. S1C). Activation by this mutant is similar to wildtype *Gα*_*q*_ (21-24), and migration on a size exclusion column as a complex with *PLCβ3* is indistinguishable from wildtype *Gα*_*q*_ (Fig. S1A-B). When *Gα*_*q*_ is added to the enzyme assay in the absence of *PLCβ3*, it does not affect GIRK current (Fig. S1D).

Fig. 2A shows the influence of *Gα*_*q*_ on *PIP2* hydrolysis. In the presence of 29 *nM PLCβ3* in aqueous solution the decay of GIRK-ALFA current is slow (reduction of ~15% over 30 seconds), reflecting slow hydrolysis of *PIP2*. In the presence of 1.0 *µM Gα*_*q*_, by contrast, the current reduction is faster, reflecting more rapid hydrolysis. The red curves correspond to fits to Eq. 8 and yield *V*_*max*_=0.0031 *mol%/s, K*_*M*_=0.52 *mol%* (R^2^=0.97) in the absence of *Gα*_*q*_ and *V*_*max*_=0.091 *mol%/s, K*_*M*_=0.42 *mol%* (R^2^=0.99) in the presence of 1.0 *µM Gα*_*q*._ By performing these experiments in multiples, with different concentrations of *Gα*_*q*_ (0 to 1000 *nM*) in the aqueous solution that interfaces the lipid bilayer, we observe that *Gα*_*q*_ increases *V*_*max*_ without affecting *K*_*M*_ (Fig. 2B-C, S2A-F). The red dashed curve in Fig. 2B is a rectangular hyperbola with a half activation concentration for *Gα*_*q*_ about 120 nM. The maximum increase in *V*_*max*_ elicited by *Gα*_*q*_ is about 35-fold above *V*_*max*_ in the absence of *Gα*_*q*_. Previous work showed a 20 to 50-fold enhancement of PLC*β3* catalysis, but any relationship to *V*_*max*_ or *K*_*M*_ was unknown in earlier studies of *Gα*_*q*_ (9-11). We note that the effect of *Gα*_*q*_ to increase *V*_*max*_ without affecting *K*_*M*_ is exactly what we observed for *Gβγ* stimulation of *PLCβ3* (14). However, as we will show below, the origins of these apparently similar effects are mechanistically distinct.

### Gα_q_ modifies k_cat_ of PLCβ3 catalysis

Because *PLCβ3* is an aqueous soluble enzyme that must partition onto the membrane surface to catalyze *PIP2* hydrolysis, the kinetic parameter *V*_*max*_ is the product of two separate quantities, expressible as

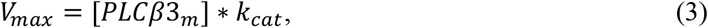

where [*PLCβ3*_*m*_] is the local concentration of *PLCβ3* on the membrane surface (subscript m) and *k*_*cat*_ is the first order rate constant for the hydrolysis of *PIP2* by a *PLCβ3-PIP2* complex on the membrane surface. In principle, to increase *V*_*max*_, *Gα* could affect either or both quantities. To examine whether *Gα*_*q*_ affects the membrane concentration of *PLCβ3*, we measured whether *Gα*_*q*_ changes the degree to which *PLCβ3* partitions onto the membrane, i.e., whether *Gα* recruits *PLCβ3* to the membrane surface. The concentration of *PLCβ3* at the membrane is determined by its partition coefficient, *K*_*x*_, which is the ratio of the mole fraction *PLCβ3* on the membrane [*PLCβ*_*m*_] to the mole fraction of *PLCβ3* in aqueous solution [*PLCβ*_*w*_]:

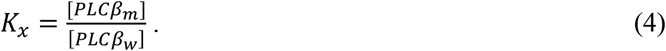

We used a vesicle spin down assay to measure the fraction of *Gα* or *PLCβ3* in the absence and presence of *Gα*_*q*_ that binds to the membrane (*F*_*p*_). This was done by incubating large unilamellar vesicles (LUVs) consisting of 2DOPE:1POPC:1POPS (wt:wt:wt) with *Gα*_*q*_ or *PLCβ3* (*± Gα*_*q*_), centrifuging the mixture, and then measuring the fraction of *Gα*_*q*_ or *PLCβ3* associated with the membranes (Fig. S4A-C). These experiments were carried out at several lipid concentrations and the measured *F*_*p*_ for *Gα* or *PLCβ3* (*± Gα*) was graphed as a function of lipid concentration (Fig. 2D, S4E-G). The dashed curves correspond to the function

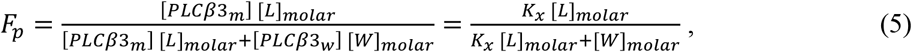

where [*W*]_*molar*_ is the molar concentration of water, ~55 *M*, and [L]_molar_ is half the total lipid concentration, recognizing that proteins can only access the outer leaflet of the LUVs. Therefore, *K*_*x*_ is the only free parameter (Fig. 2D, S4F-G) (14, 25). The results show that neither wildtype nor Q209L mutant *Gα*_*q*_ affect partitioning of *PLCβ3* onto these membrane surfaces (Fig. 2D, S4B-C, F). Moreover, *Gα*_*q*_ alone, at least the soluble version used in these experiments, does not partition onto membranes in our experiments, in contrast to previously reported results (15) (Fig. S4A, E).

Having established that *Gα*_*q*_ in these studies does not increase the membrane-bound concentration of *PLCβ3*, from Eq. 3 we conclude that the increase in *V*_*max*_ in the presence of *Gα*_*q*_ must come from an increased *k*_*cat*_. In the experiments shown in Fig. 2B, the aqueous concentration of *PLCβ3* was 29 *nM* = 5.×10^−8^ *mol%*, which, using the partition coefficient *K*_*x*_ = 2.9×10^4^ (14) and Eq. 4, yields a membrane concentration for *PLCβ3*, [*PLCβ*_*m*_] = 1.5×10^−3^ *mol%*. Therefore, from *V*_*max*_ = 0.091 *mol%/s* (Fig. 2A) and Eq. 3, we calculate *k*_*cat*_ ~60 s^−1^ in the presence of a maximally-activating concentration of *Gα*_*q*_, which is about 35-fold higher than *k*_*cat*_ in the absence of *Gα*_*q*_ (Fig. 2A, Table 1) (14). This finding contrasts the influence of *Gβγ* on *PLCβ3* function, which increases *V*_*max*_ almost entirely through membrane recruitment with little effect on *k*_*cat*_ (Table 1) (14).

**Table 1:** Kinetic parameters for *PLCβ3*.

| Condition | $K_M$ (mol%) | $k_{cat}$ ($s^{-1}$) | Fold-increase<sup>1</sup> |
| --- | --- | --- | --- |
| *PLCβ3* alone<sup>2</sup> | 0.43±0.05 | 1.7±0.5 | - |
| *PLCβ3* + $G\alpha_q$<sup>3</sup> | 0.51±0.04 | 56.9±8 | 34 |
| *PLCβ3* ΔX-Y all | 0.33±0.04 | 1,977.5±150 | 1,160 |
| *PLCβ3* ΔX-Y all + $G\alpha_q$<sup>4</sup> | 0.31±0.02 | 2,485±250 | 1,460 |
| *PLCβ3* ΔX-Y contact | 0.36±0.02 | 588.5±90 | 346 |
| *PLCβ3* ΔX-Y contact + $G\alpha_q$<sup>4</sup> | 0.34±0.04 | 1,161.3±140 | 679 |

1. Over wildtype basal activity 2. Previously determined (14) 3. For saturating $G\alpha_q$ , 300 nM 4. For 200 nM $G\alpha_q$

Our observation that *Gα*_*q*_ increases *k*_*cat*_ without discernably affecting *K*_*M*_ places constraints on the rate constants of a Michaelis-Menten kinetic reaction scheme. Specifically, for the reaction 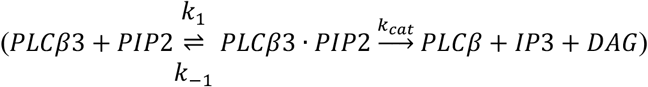, where *k*_*1*_ and *k*_*-1*_ are the forward and reverse rate constants for *PLCβ3-PIP2* complex formation and *k*_*cat*_ the catalytic step, *K*_*M*_ is

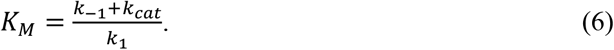

The most likely explanation for a 35-fold change in *k*_*cat*_ with little effect on *K*_*M*_ is that *k*_*-1*_ >> *k*_*cat*_ so that the value of the numerator is little affected by changes in the smaller quantity, *k*_*cat*_. In the framework of the above reaction scheme, this would imply that most encounters between *PLCβ3* and *PIP2* dissociate prior to hydrolysis.

### Gα_q_-dependent activation is dependent on the X-Y linker

A natural explanation for how *Gα*_*q*_ increases *k*_*cat*_ is that it somehow destabilizes the interaction between the autoinhibitory X-Y linker and the active site, allowing it to be displaced with a higher probability. To test this possibility, we expressed and purified *PLCβ3* lacking the entire X-Y linker (R471-V584, PLC*β*3 *β*X-Y all) or the segment of the linker in direct contact with the active site (T575-V584, *Δ*X-Y contact) and tested their basal and *Gα*_*q*_-dependent catalytic activity. If *Gα*_*q*_-dependent activation is mediated through the X-Y linker, then the maximum fold-activation by *Gα*_*q*_ should be significantly reduced, which has been previously reported (13, 26).

Both X-Y linker mutants exhibited significantly increased basal (i.e., unstimulated by G proteins) *V*_*max*_: ~2,300-fold for *PLCβ3 β*X-Y all and ~700-fold for *PLCβ3 β*X-Y contact, consistent with the autoinhibitory function of the linker (Fig. 3A, S2G). Membrane partitioning experiments showed that membrane association is enhanced only ~2-fold in the *β*X-Y all construct (Fig. S4D, G). Therefore, the increase in basal activity is primarily due to an increase in *k*_*cat*_, ~1,100-fold for *PLCβ3 β*X-Y all and ~350-fold for *PLCβ3 β*X-Y contact (Table 1). This observation also indicates that partitioning is not significantly influenced by the X-Y linker. In addition, the *K*_*M*_ values for the deletion mutants were not significantly different than wildtype (Table 1), suggesting that the linker does not simply act as a competitive inhibitor, blocking *PIP2* from binding to the active site. The small difference in basal activity between the two constructs, ~3-fold, suggests that most of the autoinhibitory impact is mediated by the residues in direct contact with the active site.

**Figure 3:**
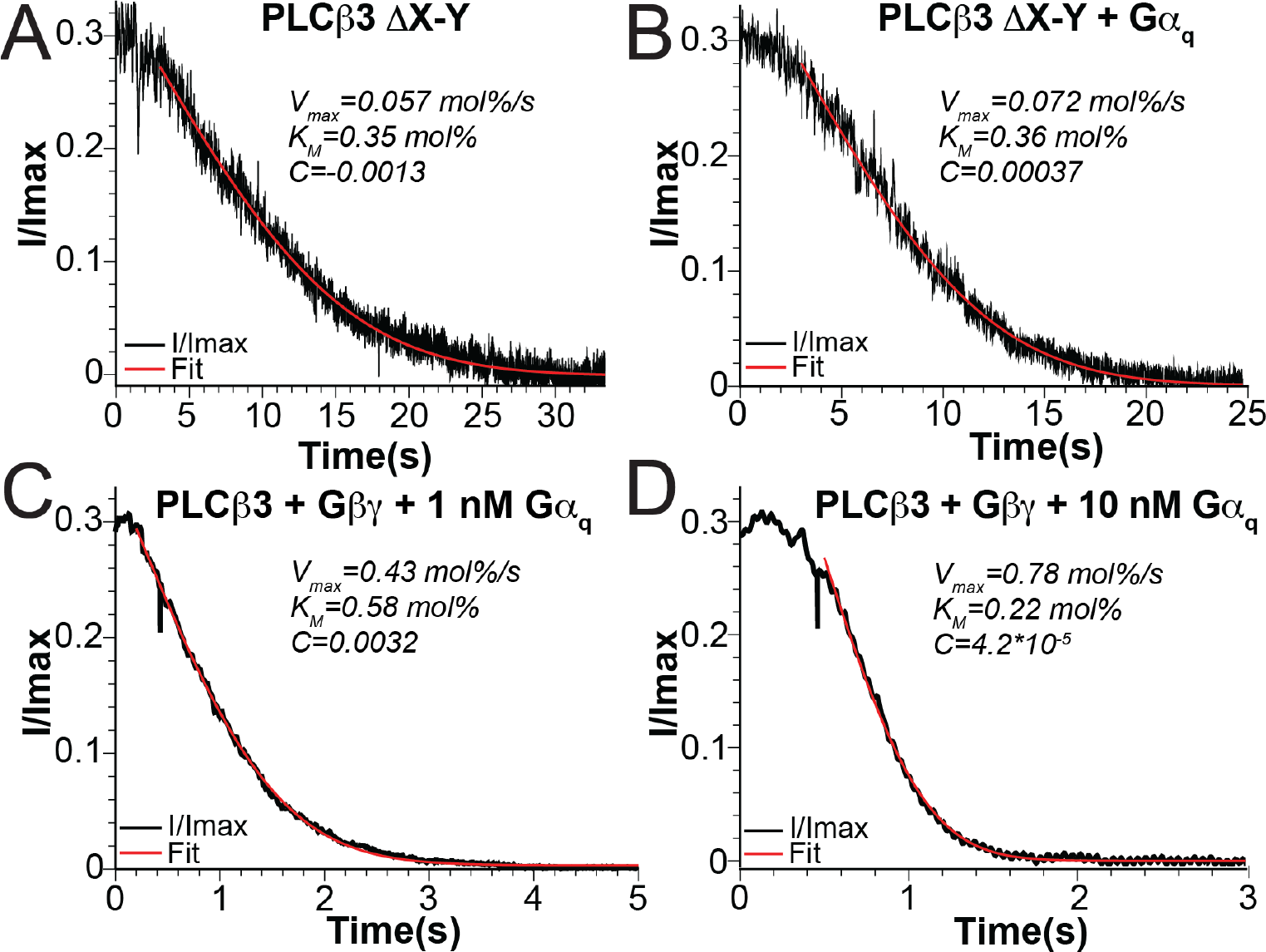
Involvement of the X-Y linker in *Gα*_*q*_-dependent activation and demonstration of dual activation of *PLCβ3* by *Gα*_*q*_ and *Gβγ*. A-B: Representative normalized current decay for *PLCβ3* lacking the entire X-Y linker (*Δ*X-Y all) using 290 *pM* of enzyme in the absence (A) or presence (B) of 200 *nM Gα*_*q*_ fit to Eq. 8 to determine *V*_*max*_ and *K*_*M*_ (red curves), R^2^=0.96, R^2^=0.98 for A and B. C-D: Representative *PLCβ3*-dependent normalized current decay in the presence of lipidated *Gβγ* and 1.0 *nM Gα*_*q*_ (C) or 10 *nM Gα*_*q*_ (D) fit to Eq. 8 (red curve) to determine *V*_*max*_ and *K*_*M*_, R^2^=1.0 for C and D.

Addition of 200 *nM Gα*_*q*_, which produces a ~20-fold increased *V*_*max*_ in wildtype *PLCβ3* (Fig. 2B), had less than a two-fold effect on *V*_*max*_ for *PLCβ3 Δ*X-Y and ~2-fold for *PLCβ3 Δ*X-Y contact (Fig. 3B, S2H, Table 1). Thus, an intact autoinhibitory X-Y linker is required for *Gα*_*q*_-dependent activation. Because the lack of *Gα*_*q*_-dependent activation is comparable in the two mutants, stimulation by *Gα*_*q*_ is likely mediated through the residues that directly contact the active site. Taken together, these results suggest that the presence of the X-Y linker in the active site is a major suppressor of *k*_*cat*_ and that *Gα*_*q*_-dependent activation is mediated through partial relief of this suppression.

One might wonder whether the relative insensitivity of the catalytic rate to *Gα*_*q*_ in the *Δ*X-Y mutants could reflect *PIP2* depletion near the active site owing to the relatively high catalytic rates in these mutants, i.e., substrate access becomes diffusion-limited. Based on a calculation presented in Appendix 2 of Supplementary Information, we think this is unlikely to be the case. More likely, allosteric regulation of the active site of *PLCβ3* is mediated at least in part through the inhibitory X-Y linker and manifests kinetically through the altered *k*_*cat*_ that we observe.

### Simultaneous activation of PLCβ3 by Gα_q_ and Gβγ

We have demonstrated that non-lipidated *Gα*_*q*_ and lipidated *Gβγ* activate *PLCβ3* through different mechanisms, *Gβγ* through membrane recruitment to increase the membrane concentration of enzyme and *Gα*_*q*_ by increasing the catalytic rate constant. Given these observations, we suspected that dual activation of *PLCβ3* by both G proteins would combine both mechanisms, which would lead to a product, rather than a sum, of the two effects (Eq. 3). To test this idea, we measured *PLCβ3* activity in the presence of a high concentration of lipidated *Gβγ* and 1.0 *nM* or 10 *nM Gα*_*q*_ (Fig. 3C-D). The current decays in the bilayer enzyme assay were very rapid, but still well-fit by the transformed Lambert W Function, thus permitting determination of *V*_*max*_ and *K*_*M*_ (Fig. 3C-D). Fig. 3C-D show that *Gα*_*q*_ induces a concentration-dependent increase in *V*_*max*_ in the presence of *Gβγ*, as was observed in the absence of *Gβγ* (Table 2). Moreover, the fold-increase of *V*_*max*_ in the presence of *Gα*_*q*_ compared to that in the absence of *Gα*_*q*_ is approximately the same whether *Gβγ* is present or not (Table 2). This supports the independent action of *Gα*_*q*_ and *Gβγ* and the conclusion that together both G proteins increase *V*_*max*_ by a produce rule.

**Table 2:** Effect of dual Activation with *Gα*_*q*_ and *Gβγ* on *V*_*max*_.

| Condition | $V_{max}$ (mol%/s) | Total fold-increase | Fold-increase over 0 $G\alpha_q$ |
| --- | --- | --- | --- |
| $PLC\beta 3$ alone<sup>1</sup> | 0.0026±0.0007 | - | - |
| $PLC\beta 3 + 1.0 \text{ nM } G\alpha_q$ | 0.0068±0.0007 | 2.6±0.3 | 2.6 ± 0.3 |
| $PLC\beta 3 + 10 \text{ nM } G\alpha_q$ | 0.0076±0.001 | 2.9±0.4 | 2.9±0.4 |
| $PLC\beta 3 + G\beta\gamma$<sup>1</sup> | 0.17±0.02 | 65 | - |
| $PLC\beta 3 + G\beta\gamma + 1.0 \text{ nM } G\alpha_q$ | 0.34±0.1 | 129±38 | 1.8±0.5 |
| $PLC\beta 3 + G\beta\gamma + 10 \text{ nM } G\alpha_q$ | 0.55±0.1 | 213±44 | 2.9±0.6 |

1. Previously reported (14)

### Structure of the PLCβ-Gα_q_ complex on lipid vesicles

We determined the structure of the *PLCβ3-Gα*_*q*_ complex associated with lipid vesicles at 3.4 Å (Fig. 4, S5, Table S1). The sample was prepared by combining *PLCβ3* and wildtype *Gα*_*q*_ bound to GDP-AlF_4,_ purifying the complex using size exclusion chromatography (Fig. S1A), and then mixing the purified complex with lipid vesicles comprised of 2DOPE:1POPC:1POPS. The structure of the complex contains density for the *PLCβ3* catalytic core and the proximal CTD, but the CTD linker and the distal CTD are disordered (Fig. 4A-B), suggesting conformational heterogeneity of the domains with respect to each other. The overall complex is very similar to the previously determined crystal structure, including the X-Y linker engaged in the active site, with a C*α* rmsd of 0.84 Å (10) (Fig. 4B-C). Despite the disordered distal CTD, which was previously shown to be part of the *PLCβ3-Gα*_*q*_ interface (10), the interface between the *PLCβ3* catalytic core and *Gα*_*q*_ is extensive, burying ~1500 Å^2^ and involving 56 residues, 27 from *Gα*_*q*_ and 29 from *PLCβ3* (Fig. 4E, S7A, Table S2-3). Compared to the structure of the catalytic core in the absence of *Gα*_*q*_, the only conformational difference is the displacement of the H*α*2’ away from the catalytic core (Fig. 4D). Despite its proximity to the catalytic site, the H*α*2’ displacement does not induce additional changes in that region (Fig. 4D). Membrane association of the complex also does not produce conformational differences other than the additional heterogeneity between the catalytic core and the distal CTD (Fig. 4C).

**Figure 4:**
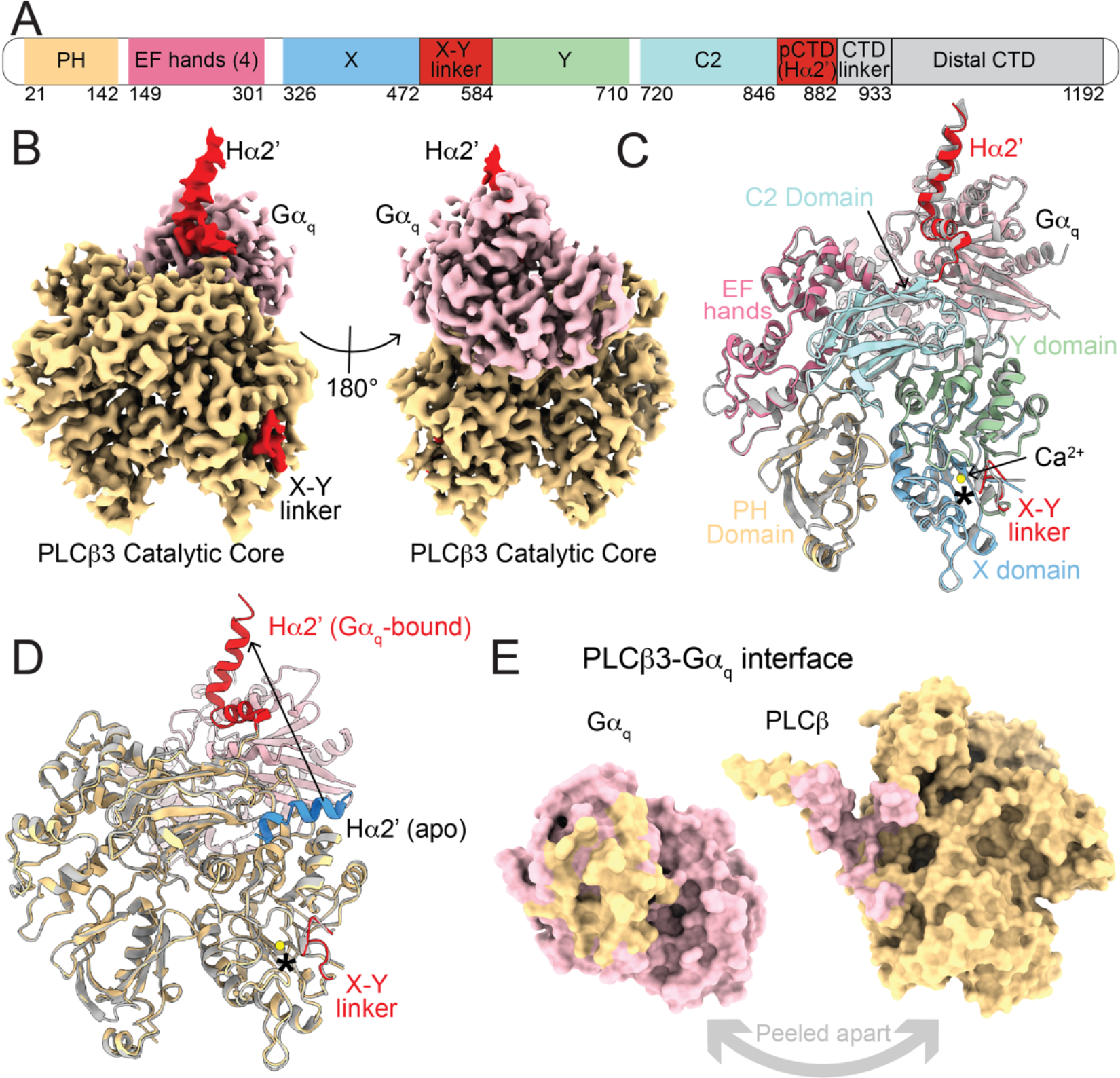
Structure of the *PLCβ3-Gα*_*q*_ complex on lipid vesicles. A: Primary structure arrangement of *PLCβ3* enzymes. Sections are colored by domain. The PH domain is yellow, the EF hand repeats are pink, the C2 domain is light teal, the Y domain is green, the X domain is light blue, the X-Y linker and the pCTD are red. Domains in gray (CTD linker and Distal CTD) are not observed in our structures. pCTD is proximal CTD. B: Sharpened, masked map of the *PLCβ3-Gα*_*q*_ complex colored by protein. *PLCβ3* is yellow and *Gα*_*q*_ is pink. The autoinhibitory elements in *PLCβ3*, the X-Y linker and the pCTD, are colored red. C: Structural alignment of the *PLCβ3-Gα*_*q*_ complex on membranes, colored by domain as in A, with the previously determined crystal structure of the complex (PDBID: 4GNK, (10)), in gray. C*α* rmsd is 0.84 Å. Calcium ion from the cryo-EM structure is shown as a yellow sphere and the active site is denoted with an asterisk. D: Structural alignment of *PLCβ3-Gα*_*q*_ complex on membranes, colored by protein-*PLCβ3* is yellow and *Gα*_*q*_ is pink, with the previously determined cryo-EM structure of the apo catalytic core (PDBID: 8EMV, (14)) colored in gray. The X-Y linker and the pCTD from the *Gα*_*q*_ complex are colored red, the calcium ion from the cryo-EM structure is shown as a yellow sphere and the active site is denoted with an asterisk. The H*α*2’ from the apo structure is colored in blue to highlight its position on the catalytic core and an arrow denotes the *Gα*_*q*_-dependent movement of the H*α*2’. E: Surface representation of the *PLCβ3-Gα*_*q*_ interface peeled apart to show extensive interactions. Residues on *PLCβ3* that interact with *Gα*_*q*_ are colored in pink and residues on *Gα*_*q*_ that interact with *PLCβ3* are colored in yellow. Interface residues were determined using the ChimeraX interface feature using a buried surface area cutoff of 15 Å^2^.

Despite our finding that the X-Y linker is involved functionally in *Gα*_*q*_-dependent activation, we do not observe structural differences at the active site or its interface with the X-Y linker, even with extensive classification targeting that region. This observation is not too surprising, however, given the magnitude of activation of *Gα*_*q*_ compared to the activity in the absence of linker. The basal *k*_*cat*_ and maximal *Gα*_*q*_-stimulated *k*_*cat*_ are only ~0.09% and ~3%, respectively, of the activity in the absence of the linker, suggesting that *Gα*_*q*_ does not alter the probability of its occupancy in the active site enough to be observable in structural experiments. In other words, if we take the activity in the absence of the linker as zero occupancy of the linker in the active site, then even in the presence of saturating *Gα*_*q*_, the linker would only be displaced 3% of the time, which is not easily detectable using cryo-EM.

### Structure of the PLCβ-Gβγ(2)-Gα_q_ complex on lipid vesicles

We also determined the structure of the *PLCβ3-Gβγ(2)-Gα*_*q*_ complex bound to lipid vesicles to 3.4 Å resolution (Fig. 5, S6, Table S1). We reconstituted lipidated *Gβγ* into vesicles comprised of 2DOPE:1POPC:1POPS as previously described (14), mixed *PLCβ3* and wildtype *Gα*_*q*_ bound to GDP-AlF_4_ and added the complex to the *Gβγ*-containing lipid vesicles for grid preparation. The structure contains the *PLCβ3* catalytic core and proximal CTD, two *Gβγ* molecules, and one *Gα*_*q*_ molecule (Fig. 5, S6). The CTD linker and distal CTD were disordered, as in the other structures of *PLCβ3*-G protein complexes on membranes (14). The structure is very similar to the structures of *PLCβ3* in complex with each G protein on its own, with no additional conformational changes observed (Fig. 5B-C). The X-Y linker is present in the active site and the H*α*2’ is in the *Gα*_*q*_-bound conformation (Fig. 5A-B). Each of the *PLCβ3*-G protein interfaces is unaltered by the presence of the additional G protein (Fig. 5, S7B-D, Table S2-S3). These observations are consistent with the functional experiments, which show that binding of one G protein does not influence the other, and that they act independently to give a product rule for catalytic enhancement when both G proteins are present.

**Figure 5:**
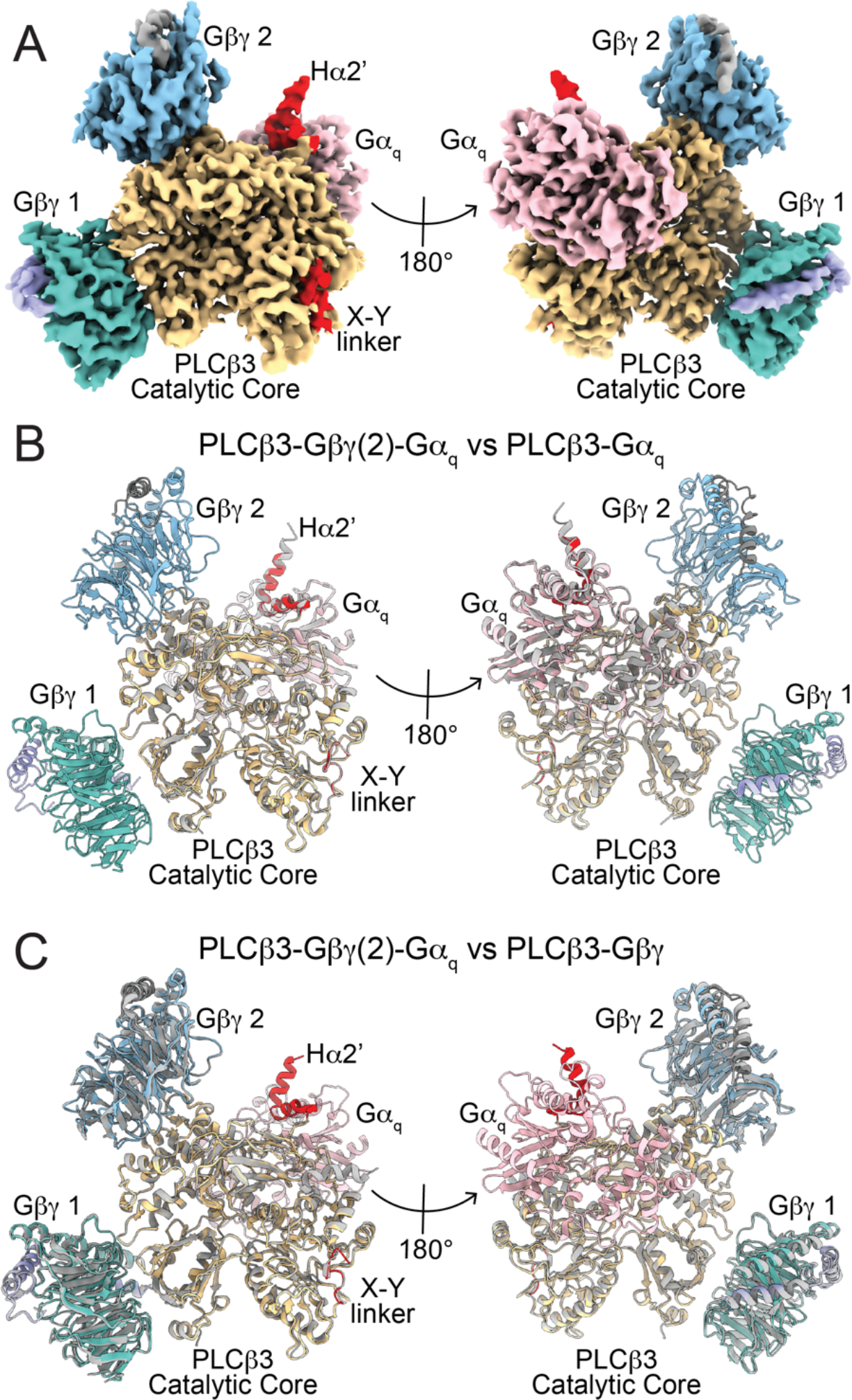
Structure of the *PLCβ3-Gβγ(2)-Gα*_*q*_ complex on lipid vesicles. A: Sharpened, masked map of the *PLCβ3-Gβγ(2)-Gα*_*q*_ complex colored by protein. *PLCβ3* is yellow, *Gα*_*q*_ is pink, *Gβ* 1 is dark teal, *Gγ* 1 is purple, *Gβ* 2 is light blue, *Gγ* 2 is gray. The X-Y linker and pCTD are colored red. B-C: Structural alignment of *PLCβ3-Gβγ(2)-Gα*_*q*_ complex on lipid vesicles, colored as in A, with the *PLCβ3-Gα*_*q*_ complex on lipid vesicles in gray (B, C*α* rmsd=0.62 Å) or with the *PLCβ3-Gβγ* complex on lipid vesicles in gray (C, C*α* rmsd=0.63 Å), (PDBID: 8EMW, (14)).

### Membrane association of PLCβ3-G protein complexes

Unmasked classification on the aligned particle subsets for each complex yielded reconstructions with density for the lipid bilayer, allowing us to study the orientation of each complex on the membrane (Fig. 6). Two different membrane-associated reconstructions of the *PLCβ3-Gα*_*q*_ complex were observed, in which the catalytic core associates with the membrane and orients the active site towards the membrane (Fig. 6B-C). There were no differences in the protein components of each reconstruction, suggesting that the complex tilts on the membrane as a rigid body (Fig. 6B-C). This orientation differs from *PLCβ3* in the absence of G proteins, where the catalytic core extends away from the membrane (Fig. 6A) (14). This orientation also differs from the *PLCβ3-Gβγ* complex where the two *Gβγs* anchor the catalytic core to the membrane on the opposing side, resulting in the catalytic site tilting away from the membrane (14).

**Figure 6:**
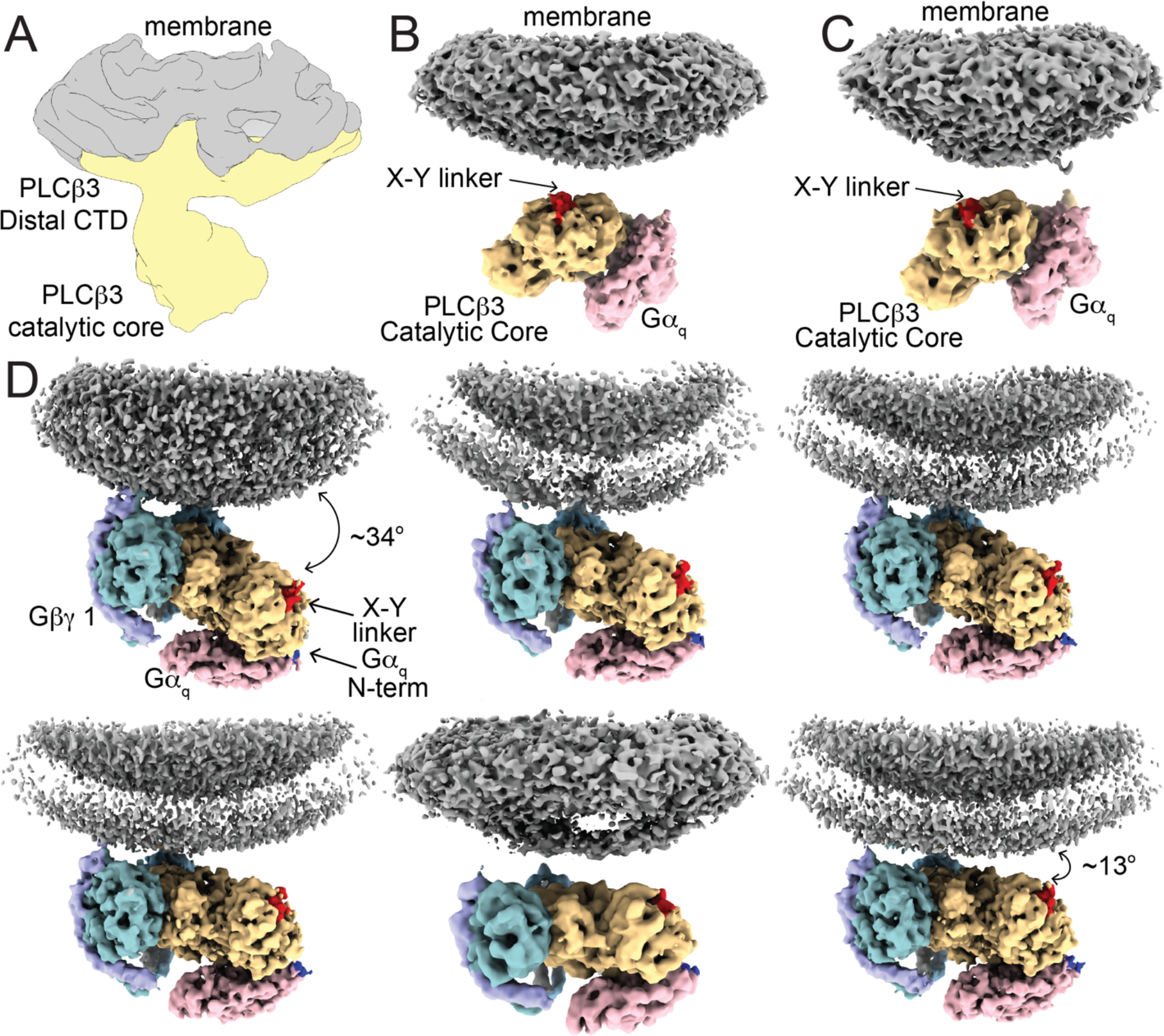
Membrane association of *PLCβ3* in the presence of G proteins. A: Unsharpened reconstruction of *PLCβ3* bound to lipid vesicles in the absence of G proteins shown for comparison (14). *PLCβ3* is colored in yellow and the membrane is colored in gray. B-C: 3D reconstructions of two different orientations of the *PLCβ3-Gα*_*q*_ complex on the membrane surface. The reconstructions are colored by protein, *PLCβ3* is yellow, *Gα*_*q*_ is pink, and the membrane is gray. The *PLCβ3* X-Y linker is colored red to highlight he active site within the catalytic core. D: 3D reconstructions of six 3D classes of the *PLCβ3-Gβγ(2)-Gα*_*q*_ complex on membranes showing different positions of the complex with respect to the membrane arranged by degree of tilting. The reconstructions are colored by protein as in B and C, and *Gβ* 1 is dark teal, *Gγ* 1 is purple, *Gβ* 2 is light blue, *Gγ* 2 is gray. The N-terminus of *Gα*_*q*_ is colored blue for reference.

The *PLCβ3-Gα*_*q*_ orientation seems more poised for catalysis as the active site is oriented directly towards the membrane (Fig. 6B). It appears likely that this orientation is driven by the *Gα*_*q*_-induced conformational change of the H*α*2’ because, when *Gα*_*q*_ binds, the H*α*2’ is displaced from the catalytic core and an underlying hydrophobic patch is exposed on the surface of the catalytic core (Fig. S8A-B). The point of membrane association in the complex is very close to this hydrophobic patch, suggesting that it plays a role in positioning the complex on the membrane (Fig. S8C-D). If we fit the catalytic core in the absence of G proteins into the density of the *PLCβ3-Gα*_*q*_ complex on the membrane, the H*α*2’ protrudes near the membrane density, suggesting that it could hinder membrane association in this configuration. This suggests that *Gα*_*q*_ binding to *PLCβ3*, even without the lipid anchor, could indirectly play a role in orienting the *PLCβ3* catalytic core on the membrane. Such an orientation effect would apply to *PLCβ3s* that have partitioned onto the membrane, rather than on the partitioning step, consistent with the observation that *Gα*_*q*_ does not alter membrane association. It is possible that this orientation of the *PLCβ3-Gα*_*q*_ complex on the membrane contributes to the *Gα*_*q*_-mediated displacement of the X-Y linker from the active site.

For the complex with both G proteins, the orientation resembles that of the *PLCβ3-Gβγα*complex, where the two *Gβγs* firmly anchor the PH domain and EF hands to the membrane and the other side of the catalytic core tilts away (Fig. 6D) (14). We also observed tilting of the complex with *PLCβ3* and both G proteins on the membrane, as in the *PLCβ3-Gβγ* complex. Six reconstructions with at least 4 Å resolution were observed with differing tilt angles of the catalytic core with respect to the membrane, ranging from 34*°* in the most tilted to 13*°* in the least tilted (Fig. 6D). There are no changes to the protein components in these reconstructions, suggesting again that the complex tilts on the membrane as a rigid body. The membrane orientation seems to be driven by the *Gβγs* under these conditions, which we speculate is due to the lipid anchor on *Gβγ* and the lack thereof on *Gα*_*q*_.

However, the observed orientations are not incompatible with a lipid anchor on *Gα*_*q*_, which would likely be present in a cell. In our structures, the N-terminus of *Gα*_*q*_ is disordered until position 38 (Fig. 6D, blue region) and the lipid modifications are placed on cysteines at positions 9 and 10. Even in our most tilted reconstruction, where the *Gα*_*q*_ N-terminus is ~85 Å from the membrane (Fig. 6D), the disordered portion is long enough for the lipid anchors to reside in the membrane. This observation is consistent with other structures of G*α* subunits in complex with their effectors, including adenylyl cyclase and TRPC5 (27-30), where the G*α* is positioned ~50 Å from the membrane and the N-terminus is disordered. These observations are consistent with our functional experiments showing that *Gα*_*q*_ activates *PLCβ3* by increasing *k*_*cat*_ rather than through membrane recruitment. But it is possible that a lipidated *Gα*_*q*_ might also recruit *PLCβ3* to the membrane in addition to increasing its *k*_*cat*_.

## Discussion

In a recent study we analyzed the structural and enzymatic properties of *PLCβ3* in the absence and presence of *Gβγ* on lipid vesicles (14). We found that *PLCβ3* catalyzes *PIP2* hydrolysis in accordance with Michaelis-Menten enzyme kinetics with a very small *k*_*cat*_ (~1.7 s^−1^), but that *Gβγ* can increase net catalysis by binding to *PLCβ3* and thus recruiting it to the membrane. It is known that *Gα*_*q*_ also increases net catalysis (8, 10, 11, 16). In this study we investigate the influence of *Gα*_*q*_ on *PLCβ3* activity. We used *Gα*_*q*_ that does not contain a lipid anchor. Our essential findings are as follows. 1) *Gα*_*q*_ increases *V*_*max*_ in a concentration-dependent manner, following a rectangular hyperbola, consistent with 1:1 binding of *Gα*_*q*_ to *PLCβ3*. The apparent equilibrium constant for binding is ~120 *nM*, and maximal activation is ~35-fold greater than the basal (i.e., in the absence of *Gα*_*q*_) catalytic rate. 2) *Gα*_*q*_ without a lipid anchor does not partition onto the membrane surface, nor does it influence the degree to which *PLCβ3* partitions onto the membrane surface. Thus, *Gα*_*q*_ without a covalent lipid anchor increases *V*_*max*_ by increasing *k*_*cat*_. 3) The ability of *Gα*_*q*_ to increase *k*_*cat*_ depends on the presence of the X-Y linker autoinhibitory element on *PLCβ3*. 4) *Gα*_*q*_ and *Gβγ* act independently to increase *V*_*max*_. Consequently, when both G proteins are applied simultaneously, the net increase in *PLCβ3* catalytic activity is given by the product of the two individual effects. Under the conditions in which we have studied *PLCβ3* enzyme activity, maximal dual stimulation can increase *PIP2* hydrolysis greater than 2000-fold. 5) Structures of *PLCβ3* on lipid membrane vesicles alone, with *Gα*_*q*_, with *Gβγ*, and with both G proteins together, show that two *Gβγ* and one *Gα*_*q*_ bind to *PLCβ3* simultaneously and independently, consistent with their influence on *PLCβ3* catalysis. In summary, two *Gβγ* localize (i.e., recruit) *PLCβ3* to the membrane. Independently, *Gα*_*q*_ increases *k*_*cat*_. Mutational studies support the hypothesis that *Gα*_*q*_ regulates *k*_*cat*_ allosterically through the autoinhibitory X-Y linker (Fig. 7).

**Figure 7:**
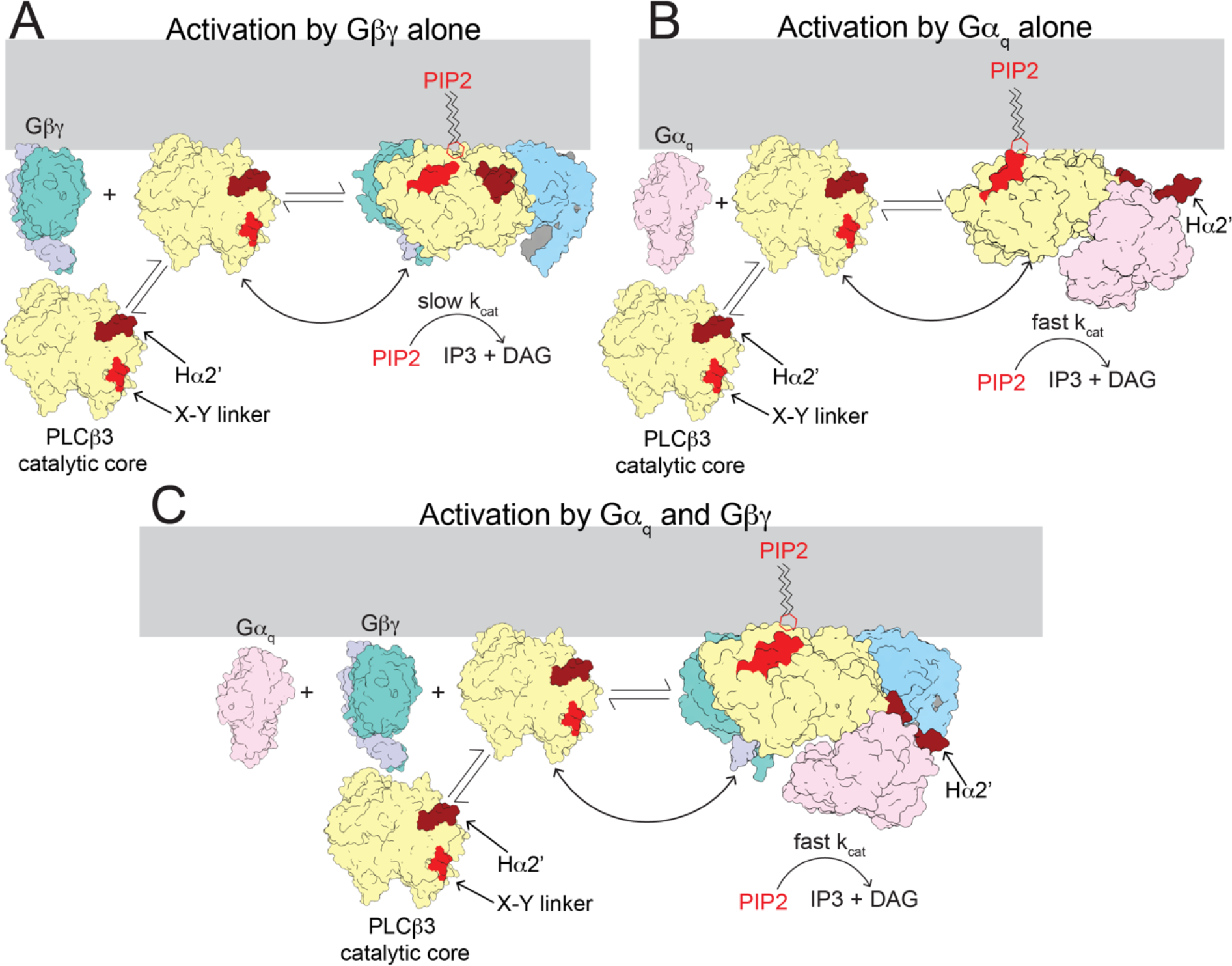
Hypothesized mechanism of activation of *PLCβ* enzymes by *Gβγ* and *Gα*_*q*_. When *PLCβ* binds the membrane, the active site is positioned away from the membrane and the enzyme is autoinhibited by both the X-Y linker and the H*α*2’, resulting in low activity in the absence in G proteins. A: Free *Gβγ* binds to membrane-associated *PLCβ*, increases its concentration at the membrane and orients the active site for catalysis, leading to an increase in *PIP2* degradation. However, the *k*_*cat*_ is limited by both the X-Y linker and the H*α*2’ (shown in red and dark red, respectively). B: Free *Gα*_*q*_ binds to membrane associated *PLCβ3*, displaces the autoinhibitory H*α*2’ (shown in dark red) and the X-Y linker is more frequently absent from the active site, resulting in an increase in *k*_*cat*_ and *PIP2* turnover. C: Free *Gα*_*q*_ and *Gβγ* simultaneously bind to membrane-associated *PLCβ*, leading to a combination of the activation effects of each G protein. The final result is increased *PLCβ3* on the membrane surface with reduced autoinhibition (both the H*α*2’ and the X-Y linker) at the membrane, leading to robust *PIP2* hydrolysis. The distal CTD of *PLCβ3* was omitted for clarity.

There is one difference in the conditions of our partitioning experiments and the kinetic experiments for *PLCβ3* function: the partitioning experiments are carried out in the absence of *PIP2*. We could not include *PIP2* in the partitioning experiments because it would be hydrolyzed throughout the measurement. However, if *PIP2* did influence the partition coefficient for *PLCβ*3, it would not affect our conclusion that *Gα*_*q*_ (without a lipid anchor) does not alter the local concentration *PLCβ*3 in the membrane, and thus increases *V*_*max*_ by increasing *k*_*cat*_. As shown in Fig. 2D, *Gα*_*q*_ does not alter the fraction of *PLCβ*3 partitioned, whereas *Gβγ* does. Enhanced partitioning caused by *Gβγ* accounts for most of its effect on catalysis (14). That *Gα*_*q*_ does not enhance partitioning is independent of the precise value of the *PLCβ*3 partition coefficient. Thus, we can attribute the ability of *Gα*_*q*_ to increase the *V*_*max*_ of *PLCβ*3 by ~35-fold as an increase in *k*_*cat*_, not its local concentration.

In the enzyme assay, *k*_*cat*_ for *PLCβ3* without *Gα*_*q*_ stimulation is ~1.7 s^−1^ (14), with maximal *Gα*_*q*_ stimulation ~60 s^−1^, and with the X-Y linker removed by mutation ~2,000 s^−1^. If we take 2,000 s^−1^ as the magnitude of *k*_*cat*_ without autoinhibition, then wildtype *PLCβ3* in the absence of *Gα*_*q*_ is inhibited by the X-Y linker more than 99.9% of the time and in the presence of a maximally-activating concentration of *Gα*_*q*_ it is still inhibited about 97% of the time. On top of this, the partition coefficient of *PLCβ3* is such that nearly all of it in a cell is in the aqueous solution, not on the membrane, in the absence of G protein stimulation (14). Why has nature so severely suppressed the catalytic activity of this enzyme? The answer, we propose, is that excessive background activity of *PLCβ3* activity will have severe consequences for the stability of cells. In fact, naturally occurring mutations show this to be the case (31-33). Not only does *PIP2* regulate the activity of many membrane channels, transporters and receptors, but of equal importance, the products of *PLCβ3*-mediated *PIP2* hydrolysis, DAG and IP3, regulate protein kinase C and the IP3 receptor, which control phosphorylation of many proteins and intracellular Ca^2+^ concentration, respectively. Therefore, we propose there is strong evolutionary ‘pressure’ to minimize baseline *PLCβ3* activity. Combining the results in our previous study (14) and in the present study we can understand how, in the setting of intense catalytic suppression, catalysis still occurs in abundance when it is called for (Fig. 7). *Gβγ*, by binding to *PLCβ3*, recruits it to the membrane (Fig. 7A). Simultaneously, *Gα*_*q*_ can increase *k*_*cat*_ ~35 fold through partial relief of X-Y linker inhibition (Fig. 7B). We show that under the conditions of our experiments, together these two regulatory mechanisms can enact a greater than 2000-fold increase in *PLCβ3* activity (Fig. 7C).

While our experiments leave little question about the involvement of the X-Y linker in *Gα*_*q*_-dependent activation, it remains unclear exactly how *Gα*_*q*_ binding alters the association of the linker in the active site. The only observed conformational change in the protein upon *Gα*_*q*_ binding is the displacement of the H*α*2’ away from the catalytic core (Fig. 4D). Perhaps the displacement of this helix increases the dynamics in the catalytic core, allowing the X-Y linker to be displaced more frequently as previously proposed (34, 35). We also observed a *Gα*_*q*_-dependent change in orientation of the catalytic core on the membrane, which could be related to the H*α*2’ displacement (Fig. 6B-C, S8). This change in membrane orientation is consistent with previous results showing that the membrane plays a role in H*α*2’ autoinhibition and that *Gα*_*q*_ only activates *PLCβ3* in the presence of membranes (13). In the *Gα*_*q*_-dependent orientation, the *PLCβ3* active site is oriented towards the membrane, which could potentially displace the X-Y linker through repulsion of its adjacent acidic stretch by the negatively charged lipids. Such a mechanism has been previously proposed (8, 13), but our observations offer a new subtlety in that the linker could be transiently displaced based on the orientation of the catalytic core on the membrane rather than a stable displacement following membrane partitioning. Involvement of the H*α*2’, as in either of these potential mechanisms, leads to the proposal that the autoinhibitory function of the H*α*2’ is related to its coupling to the X-Y linker. However, previous studies have proposed that the autoinhibition by the H*α*2’ and the X-Y linker are independent (9). Further experiments are necessary to fully understand the mechanism of X-Y linker displacement by *Gα*_*q*_ and H*α*2’ autoinhibition.

As described above, the results from our reconstitution experiments have many implications for signaling in the cellular environment. For example, the observed affinity of *PLCβ3* for *Gα*_*q*_ is relatively high, suggesting that a low level of receptor stimulation can lead to robust *PLCβ3* signaling. This effect would be further amplified in the cellular context with lipidated *Gα*_*q*_, which might also increase the local concentration of *PLCβ* on the membrane. Furthermore, because *Gβγ* and *Gα*_*q*_ activate *PLCβ3* by different mechanisms and co-activate as the product of the two fold-effects of each G protein, *PLCβ3* is well poised to serve as a coincidence detector of co-stimulation by G*α*_I_ and *Gα*_*q*_ coupled receptors, even under low levels of co-stimulation, which would be important for many physiological processes (8, 17, 18).

## Materials and Methods

### Protein Expression and Purification

Protein expression and purification was caried out as previously described (14). Following expression of all proteins, cell pellets were flash frozen and stored at −80°C until use. All protein purification steps were carried out at 4°C. Protease inhibitors (12.5 *µg/µL* leupeptin, 12.5 *µg/µL* pepstatin A, 625 *µg/µL* AEBSF, 1 *mM* Benzamadine, 100 *µg/µL* Trypisin inhibitor, 1x aprotinin, and 1 *mM* PMSF) and dNase were added to all cell lysates. GFP nanobody-coupled Sepharose resin used for protein purification was prepared as previously described (14).

The *PLCβ3* construct was comprised of human *PLCβ3* residues 10-1234 (provided by Dr. Sondek (13), downstream of GFP with a 3C protease cleavage site between the GFP and *PLCβ3*. For the X-Y linker deletion constructs, deleted residues (471-584 for *Δ*X-Y all, and 575-585 for *Δ*X-Y contact) were replaced with a 12 amino acid GSSG linker using the KLD enzyme mix (NEB). Expression and purification were carried out the same as for the wildtype protein. High Five insect cells were infected with 15-20 *mL* P3 virus per liter and harvested after 36-48 hours by centrifugation at 3,500 x *g* for 15 minutes. Protease inhibitors were used in all buffers, except the *PLCβ3* column wash buffer in which leupeptin, pepstatin A and PMSF were excluded. Cells were resuspended in *PLCβ3* lysis buffer (50 *mM* HEPES pH 8.0, 50 *mM* NaCl, 10 *mM* 2-mercaptoethanol, 5% glycerol (v/v), 0.1 *mM* EDTA, and 0.1 *mM* EGTA) and lysed by brief sonication. Lysate was clarified by centrifugation at 39,000 x *g* for 45 minutes and the supernatant was bound to GFP nanobody-coupled Sepharose resin for one hour. The resin was washed in batch with 10 column volumes of *PLCβ3* column wash buffer (20 *mM* HEPES pH 8.0, 400 *mM* NaCl, 10 *mM* 2-mercaptoethanol, 2% glycerol (v/v), 0.1 *mM* EDTA, and 0.1 *mM* EGTA), then loaded onto a column and washed with an additional 10 column volumes by gravity flow. Protein was eluted by cleavage with 3C PreScission protease for 1.5 hours and concentrated to ~10 *mg/mL* using a 15-*mL* Amicon concentrator with 100-kD molecular weight cutoff. Concentrated protein was subjected to size exclusion chromatography using a Superdex 200 10/300 increase column equilibrated with *PLCβ3* SEC buffer (20 *mM* HEPES pH 8.0, 100 *mM* NaCl, 5 *mM* Dithiothreitol (DTT), 2% glycerol (v/v), 0.1 *mM* EDTA, and 0.1 *mM* EGTA). Fractions containing *PLCβ3* were pooled, flash frozen, and stored at −80°C for later use.

To purify *PLCβ3* for non-specific cysteine labeling, the 2-mercaptoethanol was replaced with 2 *mM* tris(2-carboxyethyl)phosphine (TCEP) and peptide-based protease inhibitors were excluded from the labeling buffer. The protein-loaded resin was washed with *PLCβ3* labeling buffer (20 *mM* HEPES pH 7.4, 400 *mM* NaCl, 2 *mM* TCEP, 2% glycerol (v/v), 0.1 *mM* EDTA, and 0.1 *mM* EGTA) prior to 3C PreScission protease cleavage. Following elution, maleimide LD655 (36) was added in 5-fold molar excess and incubated overnight, protected from light. Labeled protein was concentrated to ~10 *mg/mL* using a 15-*mL* Amicon concentrator with 100-kD molecular weight cutoff and subjected to size exclusion chromatography using a Superdex 200 10/300 increase column in *PLCβ3* SEC buffer. Fractions with labeled *PLCβ3* were pooled and the labeling efficiency was evaluated. Aliquots were flash frozen and stored at −80°C for later use. Labeling efficiency was consistently 30-40%.

For lipidated *Gβγ*, untagged human *Gβ*1 was co-expressed with human *Gγ*2 with an N-terminal His-YFP tag in High Five insect cells by infection with 12 and 8 *mL* of P3 baculovirus, respectively. Cells were harvested 36-48 hours after infection by centrifugation at 3,500 x *g* for 15 minutes. Pellets were resuspended in *Gβγ* lysis buffer (25 *mM* Tris-HCl pH 8.0, and 125 *mM* NaCl) supplemented with 5 *mM* EGTA and 5 *mM* DTT and lysed by manual homogenization. Membranes were collected by centrifugation at 39,000 x *g* for 30 minutes, resuspended in fresh *Gβγ* lysis buffer supplemented with dNase and protease inhibitors and manually homogenized again. *Gβγ* was extracted using 1% sodium cholate for 1.5 hours and centrifuged at 39,000 x *g* for 30 minutes. The supernatant was bound in batch to TALON resin equilibrated with *Gβγ* lysis buffer supplemented with 1% sodium cholate for one hour. The resin was washed in batch with 10 column volumes of *Gβγ* lysis buffer supplemented with 1% sodium cholate then loaded onto a column and washed by gravity flow with 10 column volumes of high salt buffer (25 *mM* Tris-HCl pH 8.0, 500 *mM* NaCl, and 1% sodium cholate) and 10 *mM* imidazole buffer (25 *mM* Tris-HCl pH 8.0, 125 *mM* NaCl, 1% sodium cholate, and 10 *mM* imidazole). Protein was eluted with *Gβγ* lysis buffer supplemented with 1% sodium cholate and 200 *mM* imidazole and concentrated to ~2 *mL* using a 15-mL Amicon concentrator with 30-kD molecular weight cutoff. The final protein was diluted to ~20 *mL* using *Gβγ* lysis buffer supplemented with 1% sodium cholate and the His-YFP was removed by cleavage with 3C PreScission protease overnight. The free His-YFP was removed using TALON resin equilibrated with *Gβγ* lysis buffer supplemented with 1% sodium cholate and 20 *mM* imidazole and the cleaved *Gβγ* was concentrated to 1 *mL* using a 15-*mL* Amicon concentrator with 30-kD molecular weight cutoff. Protein was purified further with size exclusion chromatography using a Superdex 200 10/300 increase column in *Gβγ* lysis buffer supplemented with 1% sodium cholate and 5 *mM* DTT. Fractions containing *Gβγ* were pooled and concentrated to 5-10 mg/mL using 4-*mL* Amicon concentrator with 30-kD molecular weight cutoff and immediately used for reconstitution.

Non-lipidated *Gβγ* was generated by introducing the C68S mutation into the *Gγα*construct, which prevents lipidation ((14, 37). In this work, the YFP was retained on the final protein. Untagged human *Gβ1* was co-expressed with human *Gγ2* C68S with an N-terminal His-YFP tag in High Five cells by infection with 12 and 8 *mL* of P3 baculovirus, respectively. Cells were harvested 36-48 hours after infection by centrifugation at 3,500 x *g* for 15 minutes. Pellets were resuspended in *Gβγ* lysis buffer and lysed by brief sonication. Lysate was clarified by centrifugation for 45 minutes at 39,000 x *g* and bound in batch to TALON resin equilibrated with *Gβγ* lysis buffer. Resin was washed in batch with 10 column volumes of *Gβγ* lysis buffer, loaded onto a column and washed by gravity flow with 10 column volumes of high salt buffer (25 *mM* Tris-HCl pH 8.0 and 500 *mM* NaCl) and 10 *mM* imidazole buffer (25 *mM* Tris-HCl pH 8.0, 125 *mM* NaCl, and 10 *mM* imidazole). Protein was eluted with *Gβγ* lysis buffer supplemented with 200 *mM* imidazole and concentrated to ~10 *mg/mL* using a 15-*mL* Amicon concentrator with 30-kD molecular weight cutoff. *Gβγ-*YFP was purified further via size exclusion chromatography using a Superdex 200 10/300 increase column equilibrated with *Gβγ* lysis buffer supplemented with 5 *mM* DTT. Fractions containing *Gβγ-*YFP were pooled, flash frozen, and stored at −80°C for later use.

For ALFA-nanobody tagged *Gβγ*, the ALFA nanobody was inserted between the N-terminal His-YFP and the human *Gγ2* gene in the background of the C68S mutant. Untagged human *Gβ1* was co-expressed with the ALFA nanobody *Gγ2* construct in High Five insect cells by infection with 12 *mL* P3 baculovirus for each construct. Cells were harvested 36-48 hours after infection by centrifugation at 3,500 x *g* for 15 minutes. Cells were resuspended in *Gβγ* lysis buffer supplemented with 5 *mM* DTT and lysed by brief sonication. Lysate was clarified by centrifugation at 39,000 x *g* for 45 minutes and bound to GFP nanobody-coupled Sepharose resin equilibrated with *Gβγ* lysis buffer for one hour. The resin was washed in batch with 10 column volumes of *Gβγ* lysis buffer then loaded into a column and washed with an additional 10 column volumes by gravity flow. Protein was eluted by cleavage with 3C PreScission protease for two hours, concentrated to 1 *mL* using a 15-*mL* Amicon concentrator with 30-kD molecular weight cutoff, and further purified by size exclusion chromatography using a Superdex 200 10/300 increase column equilibrated with *Gβγ* lysis buffer supplemented with 5 *mM* DTT. Fractions with nanobody-tagged *Gβγ* were pooled, flash frozen, and stored at −80°C for later use.

Full-length mouse GIRK2 with a C-terminal ALFA peptide tag upstream of a GFP tag was expressed using HEK293S GnTI^−^ by infection with 10% (v/v) P3 virus. 10 *mM* sodium butyrate was added 12 hours after infection and the temperature was reduced to 30°C for 48 hours. Cells were harvested by centrifugation at 3,500 x *g* for 15 minutes. Cells were resuspended in GIRK lysis buffer (25 *mM* Tris-HCl pH 7.5, 150 *mM* KCl, and 2 *mM* DTT) and lysed by manual homogenization. Membranes were collected by centrifugation at 39,000 x *g* for 30 minutes, resuspended in fresh GIRK lysis buffer supplemented with protease inhibitors and dNase and manually homogenized. GIRK was extracted from membranes with 1.5% DDM/0.3% CHS for 1.5 hours and the extraction was centrifugated at 39,000 x *g* for 30 minutes. The supernatant was bound to GFP nanobody-coupled Sepharose resin equilibrated with GIRK wash buffer (20 *mM* Tris-HCl pH 7.5, 150 *mM* KCl, 2 *mM* DTT, and 0.05%/0.01% DDM/CHS) for one hour, washed in batch with 10 column volumes of GIRK wash buffer, then loaded into a column and washed with an additional 10 column volumes of GIRK wash buffer by gravity flow. Protein was eluted by cleavage with 3C PreScission protease for 1.5 hours, concentrated to ~10 *mg/mL* using a 15-*mL* Amicon concentrator with 100-kD molecular weight cutoff, and further purified by size exclusion chromatography using a Superose 6 10/300 increase column equilibrated with GIRK SEC buffer (20 *mM* Tris-HCl pH 7.5, 150 *mM* KCl, 10 *mM* DTT, and 0.025%/0.005% DDM/CHS). Fractions with GIRK were pooled, concentrated to ~2 *mg/mL* and used for reconstitution immediately.

Human *Gα*_*q*_ was truncated to the methionine at position 7, which prevents palmitoylation (15), and inserted into the pFastBac vector downstream of an 8-His tag and a 3C protease cleavage site. This construct was previously shown to activate *PLCβ* enzymes (9-11, 15). Baculovirus was generated according to the manufactures protocol (Invitrogen). *Gα*_*q*_ was co-expressed at a 2:1 ratio with untagged rat Ric8A to improve expression (38). High Five insect cells were infected with 7 *mL* of Ric8A and 15 *mL Gα*_*q*_ p3 virus per liter of culture and harvested after 36-48 hours by centrifugation at 3,500 x *g* for 15 minutes. Cells were resuspended in *Gα*_*q*_ lysis buffer (25 *mM* Tris-HCl pH 8.0, 125 *mM* NaCl, 5 *mM* MgCl_2_, 5 *mM* 2-mercaptoethanol and 10 *mM* imidazole) supplemented with 30 *µM* GDP and lysed by brief sonication. Lysate was clarified by centrifugation at 39,000 x *g* for 45 minutes and bound in batch to Ni-NTA resin equilibrated with *Gα*_*q*_ lysis buffer supplemented with 30 *µM* GDP. The resin was washed in batch with 10 column volumes of *Gα*_*q*_ lysis buffer supplemented with 30 *µM* GDP, loaded onto a column and washed by gravity flow with 10 column volumes of high salt buffer (25 *mM* Tris-HCl pH 8.0, 500 *mM* NaCl, 5 *mM* MgCl_2_, 5 *mM* 2-mercaptoethanol, 10 *mM* imidazole and 30 *µM* GDP), 25 *mM* imidazole buffer (25 *mM* Tris-HCl pH 8.0, 125 *mM* NaCl, 5 *mM* MgCl_2_, 5 *mM* 2-mercaptoethanol, 25 *mM* imidazole, and 30 *µM* GDP) and 40 *mM* imidazole buffer (25 *mM* Tris-HCl pH 8.0, 125 *mM* NaCl, 5 *mM* MgCl_2_, 5 *mM* 2-mercaptoethanol, 40 *mM* imidazole, and 30 *µM* GDP). Protein was eluted with elution buffer (25 *mM* Tris-HCl pH 8.0, 125 *mM* NaCl, 5 *mM* MgCl_2_, 5 *mM* 2-mercaptoethanol, 250 *mM* imidazole, and 30 *µM* GDP) and concentrated to ~2 *mL* using a 15-*mL* Amicon concentrator with 30-kD molecular weight cutoff and diluted to ~20 *mL* using *Gα*_*q*_ dilution buffer (25 *mM* Tris-HCl pH 8.0, 125 *mM* NaCl, 5 *mM* MgCl_2_, and 5 *mM* 2-mercaptoethanol) supplemented with 30 *µM* GDP. The His tag was cleaved by incubation with 3C PreScission protease for 1.5 hours and separated from the protein using Ni-NTA resin equilibrated with 25 *mM* imidazole buffer. Cleaved *Gα*_*q*_ was concentrated to 1 *mL* using a 15-*mL* Amicon concentrator with 30-kD molecular weight cutoff, and further purified by size exclusion chromatography using a Superdex 200 10/300 increase column equilibrated with *Gα*_*q*_ SEC buffer (25 *mM* Tris-HCl pH 8.0, 125 *mM* NaCl, 5 *mM* MgCl_2_, and 5 *mM* DTT) supplemented with 30 *µM* GDP. Fractions with *Gα*_*q*_ were pooled, flash frozen, and stored at −80°C for later use.

The Q209L mutation was generated in the same construct using the KLD enzyme mix (NEB). Expression and purification of *Gα*_*q*_ Q209L were the same as for the wildtype protein with the following modifications. High Five insect cells were infected with 4 *mL* of Ric8A and 12 *mL Gα*_*q*_ Q209L P3 virus (3:1 G*α*_q_:Ric8A) per liter of culture and harvested after ~36 hours by centrifugation at 3,500 x *g* for 15 minutes. The GDP was replaced with 100 *µM* GTP in all buffers. The GTP-bound state of the protein was confirmed after every purification using the trypsin cleavage assay described below.

For *PLCβ3-Gα*_*q*_ Q209L complex formation (Fig. S1B), *Gα*_*q*_ Q209L with GTP was mixed with *PLCβ3* in a 2:1 molar ratio in buffer containing 25 *mM* Tris-HCl pH 7.4, 125 *mM* NaCl, 3 *mM* MgCl_2_, 75 *µM* GDP, and 5 *mM* DTT and incubated on ice for one hour. The complex was purified via size exclusion chromatography using a Superdex 200 10/300 Increase column equilibrated with the same buffer.

### Trypsin cleavage assay

The nucleotide state (GTP/GDP-AlF_4_ or GDP) of *Gα*_*q*_ was evaluated using a trypsin cleavage assay as previously described (39). Purified *Gα*_*q*_ was mixed with 3 *µg* of trypsin on ice. Samples for SDS-PAGE analysis were removed after 2 minutes and 10 minutes and immediately mixed with SDS loading dye to stop the reaction. Cleavage patterns were evaluated by SDS-PAGE. The GDP-bound state of *Gα*_*q*_ produces a band at ~20 kD and the GTP/GDP-AlF_4_-bound state produces a band at ~36 kD (39)(Fig. S1C).

### Nucleotide exchange of wildtype Gα_q_

For the structure of the *PLCβ3-Gβγ(2)-Gα*_*q*_ complex and membrane partitioning studies, an additional purification step was added to ensure that all the *Gα*_*q*_ was GDP-AlF_4_-associated. *Gβγ-*YFP was bound to GFP nanobody-coupled Sepharose resin at 4°C for 2 hours to generate a *Gβγ* column and unbound *Gβγ-*YFP was removed by washing with 10 column volumes of *Gα*_*q*_ SEC buffer supplemented with 30 *µM* GDP. Purified *Gα*_*q*_ GDP was bound to the column by incubating at 4°C for 2 hours and unbound *Gα*_*q*_ was removed by washing with 10 column volumes of *Gα*_*q*_ SEC buffer supplemented with 30 *µM* GDP. *Gα*_*q*_ GDP-AlF_4_ was eluted by incubating the column with GDP-AlF_4_ buffer (25 *mM* Tris-HCl pH 8.0, 125 *mM* NaCl, 50 *mM* MgCl_2_, 10 *mM* NaF, 30 *µM* AlCl_3_, 30 *µM* GDP, and 5 *mM* DTT) at 30°C for 2-3 hours (40). The GDP-AlF_4_ state of *Gα*_*q*_ was confirmed using the trypsin cleavage assay.

### Protein Reconstitution

For all reconstitutions, lipids in chloroform were mixed and dried under a stream of argon, washed with pentane, dried under a stream of argon and incubated under vacuum overnight. GIRK and lipidated *Gβγ* were reconstituted for bilayer experiments using 3:1 ratio of 1-palmitoyl-2-oleoyl-sn-glycero-3-phosphoethanolamine (POPE): 1-palmitoyl-2-oleoyl-sn-glycero-3-phospho-(1’-rac-glycerol) (POPG). For GIRK reconstitution, lipids were resuspended in 10 *mM* K_2_HPO_4_, 450 *mM* KCl, and 10 *mM* DTT at 20 *mg/mL* and sonicated to clarity. 1% DM was added to the lipids and the mixture was sonicated again. GIRK was added to the mixture at a protein to lipid ratio of 1:10 (wt/wt), the lipid concentration was diluted to 10 *mg/mL* and incubated at 4°C for one hour. Detergent was removed at 4°C via dialysis against 10 *mM* K_2_HPO_4_ pH 7.4, 450 *mM* KCl, 10 *mM* DTT, and biobeads. *Gβγ* was reconstituted in the same way but using different buffer conditions for the lipids and dialysis (10 *mM* HEPES pH 7.4, 150 *mM* KCl, and 10 *mM* DTT), 2% sodium cholate instead of DM, and a protein to lipid ratio of 1:5 (wt:wt). In both cases, dialysis buffer was changed every 12 hours for three changes with fresh DTT added at each change. After dialysis, liposomes were incubated with ~50% volume of biobeads for five hours at 4°C, flash frozen, and stored at −80°C until use.

For structural studies and partitioning experiments, liposomes were comprised of a 2:1:1 mixture of 1,2-dioleoyl-sn-glycero-3-phosphoethanolamine (DOPE): 1-palmitoyl-2-oleoyl-glycero-3-phosphocholine (POPC): 1-palmitoyl-2-oleoyl-sn-glycero-3-phospho-L-serine (POPS). In reconstitutions for partition experiments, 0.1 *mol%* 1,2-dioleoyl-sn-glycero-3-phosphoethanolamine-N-(lissamine rhodamine B sulfonyl) (18:1 Liss Rhod PE) was included in the lipid mixture. Lipids were resuspended at 25 *mM* in reconstitution buffer (25 *mM* HEPES pH 7.4, 150 *mM* KCl, 1 *mM* MgCl_2_, and 5 *mM* DTT) and sonicated to clarity. 40 *mM* sodium cholate was added to the mixture and sonicated briefly. For protein-free liposomes for structural studies, the mixture was diluted to 20 *mM* lipid maintaining the 40 *mM* sodium cholate and incubated at 4°C for one hour. For *Gβγ-*containing liposomes, *Gβγ* was added at a protein to lipid ratio of 1:15 (wt/wt) and the lipid concentration was reduced to 20 *mM* maintaining 40 *mM* sodium cholate and incubated at 4°C for one hour. For protein-free liposomes for partitioning experiments, lipids were diluted to 10 *mM*, maintaining the 40 *mM* sodium cholate and incubated at 4°C for one hour. For all samples, detergent was removed using four exchanges of 200 *mg/mL* biobeads washed with reconstitution buffer after two hours, 12 hours, two hours, and two hours at 4°C. Liposomes for structural studies were used immediately following reconstitution to prepare grids. Liposomes for partitioning studies were flash frozen and stored at −80°C until use.

### Bilayer experiments and analysis

Experiments were carried out and analyzed as previously described (14). 2DOPE:1POPC:1POPS lipids supplemented with 1 *mol%* 1,2-dioleoyl-sn-glycero-3-phospho-(1’-myo-inositol-4’,5’-bisphosphate) (*PIP2*) were used for all experiments. Lipids in chloroform were dried under a stream of argon and resuspended at 22 *mg/mL* in decane. Two 3-*mL* chamber cups in a vertical bilayer configuration were connected by a 100 *µm* thick piece of Fluorinated ethylene propylene copolymer with a ~250 *µm* hole, which was used to paint a bilayer with the lipid-decane mixture. Bilayer buffer (25 *mM* HEPES pH 7.4, 150 *mM* KCl, 30 *mM* NaCl, 2 *mM* MgCl_2_, and 100 *µM* CaCl_2_) supplemented with 100 *µM* GTP was used in both chambers for all experiments.

The Reference electrode and the Ground electrode were connected to the Trans and Cis chambers respectively via agarose salt bridges. A magnetic stir bar was included in the Cis chamber to facilitate continuous mixing; therefore all components of the experiment were added to the Cis chamber. Voltage across the lipid bilayer was controlled with an Axopatch 200Bamplifier in whole-cell mode. The analog current signal was lowpass filtered at 1 kHz (Bessel) and digitized at 10 kHz with a Digidata 1440A digitizer. Digitized data were recorded with the software pClamp (MolecularDevices). At +80 mV, current from channels with their *PIP2*-binding sites in the Trans chamber, which is inaccessible to *PLCβ3* added to the Cis chamber, was blocked by Mg^2+^ (14). Therefore, current measured at +80 mV is only from channels sensitive to *PLCβ3*-dependent degradation of *PIP2*, allowing the current to decay completely.

*Gα*_*q*_ Q209L with GTP was used in all bilayer experiments. 30 *nM* ALFA nanobody-tagged *Gβγ* was added to the Cis chamber at the beginning of each experiment. GIRK or *Gβγ*-containing liposomes were supplemented with KCl to a final concentration of 1 *M*, sonicated briefly at room temperature, and fused to the membrane from the Cis chamber. After fusing vesicles with GIRK, baseline current at +80 mV was measured for 3-5 minutes. Then the desired concentration of *Gα*_*q*_ was added to the Cis chamber under continuous mixing and did not alter the baseline GIRK current (Fig. S1D). After at least 1 minute, 29 *nM PLCβ3* for wildtype enzyme was added to the Cis chamber under continuous mixing while recording. For experiments with wildtype *PLCβ3* and *Gβγ* and *Gα*_*q*_, *Gβγ* was incorporated into the bilayer via vesicle fusion and the *PLCβ3-Gα*_*q*_ complex was pre-formed on ice and added to the Cis chamber under continuous mixing to initiate the current decay. For experiments with *PLCβ3 β*X-Y contact or *Δ*X-Y all constructs, a final concentration of 290 *pM PLCβ3* was used. In the presence of *Gα*_*q*_, the *PLCβ3-Gα*_*q*_ complex was pre-formed on ice and added to the Cis chamber under continuous mixing to initiate the current decay. After all current decays, a voltage family was measured to ensure integrity of the bilayer and then saturating C8PIP2 (32 *µM*) was added to recover the GIRK current. Any experiment where the current did not recover was discarded. We are confident that the starting *PIP2* concentration in the membrane was not affected by vesicle fusion because the kinetics of PIP2 hydrolysis are not dependent on the number of channels that fused (14).

Analysis was carried out in Clampfit and qtiplot. To reduce the amplitude of high-frequency undesired signal owing largely to the stir bar in the recording chamber, current time series were low pass filtered at 10-20 Hz and exported to qtiplot. Decays lasting more than 100 s, i.e., at low concentrations of *Gα*_*q*_, were down sampled by a factor of 10 prior to exporting. We also analyzed our fastest decays without low pass filtering to ensure that the 10-20 Hz filter did not alter the determination of *V*_*max*_ and *K*_*M*_ to a significant degree (i.e., 10-20 Hz did not over filter the kinetic process under study). In qtipliot, decays were leak subtracted (where the baseline current remaining in the bilayer at the end of the decay, I_leak_, was subtracted from each point in the decay) and normalized to the starting GIRK current scaled according to the starting PIP2 concentration, 1.0 *mol%*, which is ~30% of the GIRK current at saturating *PIP2* concentrations (Fig. S2A).

To test the effects of DAG (Fig. S3), 1,2-Dioleoyl-rac-glycerol was mixed with bilayer lipids at 1.0 *mol%* and used to paint bilayers. This is the maximum concentration that would be present in our experiments after all the *PIP2* is hydrolyzed. To test the effects of IP3 (Fig. S3), D-myo-Inositol 1,4,5-tris-phosphate trisodium salt was solubilized in water and added to the bilayer chamber at a final concentration of 1.0 *µM*. This concentration is several orders of magnitude higher than the maximal concentration of IP3 estimated to be present in the chamber after all the *PIP2* has been hydrolyzed (~50 *pM*). The higher concentration should account for any possible higher local concentration present at the membrane following catalysis.

*K*_*M*_ and *V*_*max*_ for each experiment were determined as previously described (14). Briefly, we determined the relationship between normalized GIRK current, *I/I*_*max*_, and *PIP2* concentration in the bilayer using *PIP2* titration experiments. The relationship is described by Eq. 7,

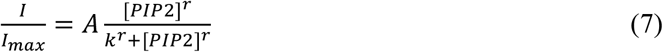

where *A*=0.88, *k*=1.5, and *r*=2.2 (14). We observed that *PLCβ3* operates according to Michaelis-Menten kinetics (Fig. 1D), described by Eq. 1 in the main text. The *PIP2* concentration as function of time was determined by integrating Eq. 1 from *τ* = 0 to *τ* = t, (Eq. 2 in the main text).

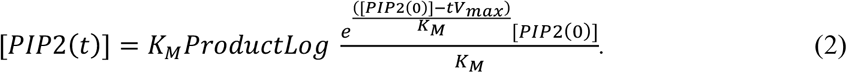

Eq. 2 was substituted into Eq. 7 and an additional term, C, was added to capture imperfections in the leak subtraction, yielding Eq. 8,

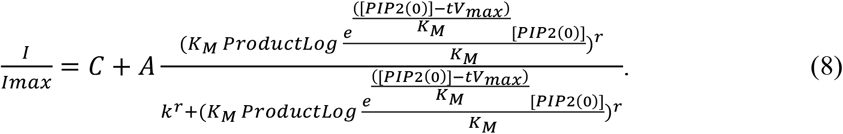

Normalized current decays were fit to Eq. 8 to determine *V*_*max*_, *K*_*M*_, and *C*, which was very small, < 1% of the normalized current.

### NMR experiments to measure lipid concentration

10-20 *µL* from each reconstitution was dissolved in ~550 *uL* of a mixture of deuterated methanol and chloroform (5:1) containing 100 *µM* of the standard sodium trimethylsilyl propionate (TSP) in 5 *mm* tubes. Proton spectra were measured on a Bruker 600 MHz instrument with a 5 *mm* HCN cryoprobe and an AVANCE NEO console. Spectra were collected at 298 K using a 30° flip angle, 16 scans, and 2.8 second acquisition time and a recycle delay of 18 *µ*s. Spectra were processed using TopSpin 4.1.1 for phasing, baseline correction and line broadening. The lipid - CH_3_ peak at 0.875 ppm was integrated relative to the TSP peak at 0 ppm and normalized to the difference in protons (9 for TSP and 6 for the lipid CH_3_) (14). The normalized peak area was used to determine the lipid concentration using the known concentration of TSP.

### PLCβ3 and Gα_q_ vesicle partition experiments

Reconstituted liposomes were subjected to 10 freeze-and-thaw cycles and extruded 21 times through a 200 *nm* membrane to produce LUVs. The reported lipid concentration is 50% of the total lipid concentration added in solution because partitioning proteins can only access the outer leaflet. Fixed concentrations of lipids were mixed with proteins of interest (wildtype *Gα*_*q*_ alone, LD655 labeled *PLCβ3 β*X-Y all alone, or wildtype LD655 labeled-*PLCβ3* and 200 *nM* wildtype *Gα*_*q*_ or 200 *nM Gα*_*q*_ Q209L) in partitioning buffer (25 *mM* HEPES pH 7.4, 150 *mM* KCl, 1 *mM* MgCl_2_, and 5 *mM* DTT) supplemented with 10 *mM* NaF, 30 *µM* GDP, and 30 *µM* AlCl_3_ for wildtype *Gα*_*q*_ or 100 *µM* GTP for *Gα*_*q*_ Q209L. Lipid-protein mixtures were incubated for one hour and centrifuged for one hour at 100,000 x *g* at room temperature. The supernatant was removed, and the membrane pellet was resuspended in an equal volume of buffer. The input, pellet, and supernatant samples were analyzed by SDS-PAGE. Samples with G*α*_q_ alone were stained using the Thermo Scientific silver stain kit and samples with *PLCβ3* were imaged using in-gel fluorescence to detect LD655-labeled *PLCβ3* (Fig. S4A-D). We previously showed that LD655-labeled *PLCβ3* has the same partition coefficient and unlabeled *PLCβ3* (14). Gel bands were quantified using Bio-Rad imagelab software. For experiments with *PLCβ3*, input, supernatant, and pellet samples were solubilized in 5% Anapoe-C12E10 to eliminate scattering artifacts and the LD655 fluorescence (ext-649, em-666) and Rhodamine fluorescence (ext-560, em-583) were measured using a Tecan plate reader. The Rhodamine signal was used to estimate the fraction of lipids that were pelleted, and the measurements were corrected for this as well as the loss of material using the difference between the input and output (pellet and supernatant) LD655 signal. Each lipid concentration was repeated with two different protein concentrations, 400 and 200 *nM* for *Gα*_*q*_ alone, 400 *nM* and 300 *nM* for *PLCβ3 β*X-Y constructs, and 200 *nM* and 100 *nM* for *PLCβ3*. Values for fraction of protein partitioned (F_p_), were determined, plotted against lipid concentration, and fit to Eq. 5 to determine *K*_*x*_ (Fig. 2D, S4E-G). For experiments with *PLCβ3*, values from the gels and the solution fluorescence measurements were consistent. The values from the solution measurements are reported.

### Cryo-EM sample preparation and data collection

For the *PLCβ3-Gα*_*q*_ complex with reconstituted liposomes, *Gα*_*q*_ was incubated on ice for 30 minutes in the presence of GDP-AlF_4_ (25 *mM* HEPES pH 7.4, 150 *mM* KCl, 5 *mM* MgCl_2_, 0.9 *mM* CaCl_2_, 10 *mM* NaF, 50 *µM* GDP, 30 *µM* AlCl_3_, and 5 *mM* DTT), then mixed with *PLCβ3* in a 2:1 molar ratio and incubated on ice for one hour. The complex was purified via size exclusion chromatography using a Superdex 200 10/300 Increase column equilibrated with 25 *mM* HEPES pH 7.4, 150 *mM* KCl, 5 *mM* MgCl_2_, 0.9 *mM* CaCl_2_, 10 *mM* NaF, 50 *µM* GDP, 30 *µM* AlCl_3_, and 5 *mM* DTT (Fig. S1A). Purified *PLCβ3-Gα*_*q*_ complex was mixed with vesicles at final concentrations of 3.6 *µM* complex and 17.5 *mM* lipids and incubated at room temperature for one hour. For the *PLCβ3-Gβγ(2)-Gα*_*q*_ complex, *Gα*_*q*_ GDP-AlF_4_ was mixed with *PLCβ3* in a 2:1 molar ratio, incubated on ice for one hour and exchanged into buffer containing 25 *mM* HEPES pH 7.4, 150 *mM* KCl, 5 *mM* MgCl_2_, 0.9 *mM* CaCl_2_, 10 *mM* NaF, 50 *µM* GDP, 30 *µM* AlCl_3_, and 5 *mM* DTT using a PD10 desalting column. Purified *PLCβ3-Gα*_*q*_ complex was mixed with *Gβγ*-containing vesicles (reconstituted at 1:15 wt/wt) at final concentrations of 4.1 *µM* complex and 16.7 *mM* lipids and incubated at room temperature for one hour.

Both samples were supplemented with 3 *mM* Fluorinated Fos-Choline-8 ~5 minutes before grid preparation. Quantifoil R1.2/1.3 400 mesh holey carbon Au grids were glow discharged for 20s, and 3.5 *µL* of sample was applied, incubated for 5 minutes at 22°C and 100% humidity, and manually blotted from below. An additional 3.5 *µL* of sample was applied and after 30 seconds of incubation, grids were blotted for 3.5s with a blot force of 0 and plunge frozen in liquid ethane using a FEI Vitrobot Mark IV. For data acquisition, grids were loaded onto a 300-kV Titan Krios transmission electron microscope, located at the HHMI Janelia Research Campus, with a Gatan K3 Summit direct electron detector and a GIF quantum energy filter with a slit width of 20 eV. 25,668 movies were collected for the *PLCβ3-Gα*_*q*_ complex and 29,832 movies were collected for the *PLCβ3-Gβγ(2)-Gα*_*q*_ complex in superresolution mode with a pixel size of 0.4195 Å and a defocus range of 1.5 to 2.5 *µm* using SerialEM (41). The movies were recorded with 50 frames, a total dose of 60 e^−^/Å^2^ (1.2 e^−^/Å^2^/frame) and a 4.26 second total exposure time (0.085s/frame) for the *PLCβ3-Gα*_*q*_ complex or a 3.86 second total exposure time (0.071s/frame) for the *PLCβ3-Gβγ(2)-Gα*_*q*_ complex.

### Cryo-EM data processing

For both complexes, motion correction was performed with 2x binning using the RELION implementation (in RELION 3.1) and CTF estimation was carried out using CTFfind4 (42-44). Particle picking was caried out using the model trained for PLC*β*3 on vesicles in crYOLO (14, 45). For the *PLCβ3-Gα*_*q*_ complex, 3,543,739 particles were picked and extracted with 2x binning and a 260 Å box size, sorted to 2,825,696 using a resolution cutoff of 4 Å, and sorted to 2,342,176 using 2D classification in cryoSPARC. Iterative rounds of *ab initio* reconstruction and heterogenous refinement in cryoSPARC were carried out to generate an initial reconstruction with density for the membrane and protein protruding (Fig. S5F). This map was used as an input for heterogenous refinement in cryoSPARC to sort particles based on membrane alignment. 592,280 particles with good membrane alignment were selected, subjected to refinement in RELION, and signal subtraction was applied to remove the membrane density. Subtracted particles were subjected to 2D classification in cryoSPARC and particles from the best 2D classes, 213,197, were used for *ab initio* reconstruction to obtain an initial map resembling the *PLCβ3-Gα*_*q*_ complex. All subtracted particles were subjected to iterative heterogenous refinement using this map as input. A final subset of 229,523 particles yielding a reconstruction with clear secondary structure features was un-subtracted and re-extracted without binning and a 289 Å box size. These particles were subjected to several cycles of Bayesian polishing and CTF refinement in RELION (46, 47) and local refinement with a mask on the entire complex in cryoSPARC. After polishing, 3D classification without alignment was carried out to improve the resolution, search for alternate configurations of the X-Y linker and obtain density for the membrane. The best subset contained 67,454 particles and resulted in a 3.4 Å reconstruction from local refinement in cryoSPARC. Two subsets containing 20,261 and 15,486 particles yielded reconstructions with density for the membrane (Fig. 6B-C). No reconstructions with differing density for the X-Y linker were observed.

For the *PLCβ3-Gβγ(2)-Gα*_*q*_ complex, 5,332,307 particles were picked and extracted with 2x binning and a 289 Å box size, sorted to 5,128,184 particles using a resolution cutoff of 5 Å, and sorted to 4,917,142 particles using 2D classification in cryoSPARC. 271,175 particles from the best 2D classes were subjected to *ab initio* reconstruction and a map with density for the membrane and two blobs of protein protruding was produced. This reconstruction was used as an input for heterogenous refinement on all 4,917,142 particles to sort based on membrane alignment. Three classes with 1,120,862, 1,235,752, and 1,682,270 particles (4,038,884 particles total) were refined separately in RELION and subjected to signal subtraction to remove the membrane density. Subtracted particles were subjected to 2D classification in cryoSPARC. 195,750 particles from the best 2D classes were subjected to *ab initio* reconstruction to obtain a map resembling the *PLCβ3-Gβγ(2)-Gα*_*q*_ complex. This map was used as an input for iterative heterogenous refinement on all 4,038,884 subtracted particles. 484,174 particles yielding a reconstruction with clear secondary structure features were un-subtracted and re-extracted without binning and a 289 Å box size. These particles were subjected to several rounds of Bayesian polishing and CTF refinement in RELION (46, 47) and local refinement with a mask on the entire complex in cryoSPARC. After polishing, 3D classification without alignment was carried out to improve the resolution, search for alternate configurations of the X-Y linker and obtain density for the membrane. The best subset contained 359,215 particles and resulted in a 3.4 Å reconstruction from local refinement in cryoSPARC. Six subsets containing 19,266, 15,138, 27,763, 21,151, 17355, and 12,194 particles yielded reconstructions with density for the membrane (Fig. 6D). No reconstructions with differing density for the X-Y linker were observed. A subset of 48,367 particles was identified that only contained density for *PLCβ3* and *Gβγ*.

Refinement of these particles yielded a reconstruction that was very similar to the previously reported *PLCβ3-Gβγ* complex and these particles were excluded from the final subset.

### Model building and Validation

For the *PLCβ3-Gα*_*q*_ complex, a previously determined crystal structure was used as the starting model (PDBID 4GNK, (10)). It was fit into the density and refined with PHENIX real-space refine (48) and manually inspected and adjusted where necessary. Regions with poor or weak density were removed. The final model consists of *PLCβ3*: 13-33, 38-92, 97-470, and 575-878 with the side chains of E14, R23, R24, M59, E60, K85, E88, E100, K169, K173, E187, R185, F197, K228, R268, K761, D777, D850, Q853, Y855, R872, R874, and Q875 truncated due poor density, and *Gα*_*q*_: 38-352 with the side chains of K98, Y103, K120, N126, D165, R166, Q197, Q265, and K276 truncated due to poor density. For the *PLCβ3-Gβγ(2)-Gα*_*q*_ complex, the *Gβγs* from the *PLCβ3-Gβγ* complex on vesicles (PDBID: 8EMW, (14) were merged with the *PLCβ3-Gα*_*q*_ complex as a starting model. It was fit into the density and refined with PHENIX real-space refine (48) and manually inspected and adjusted where necessary. Regions with poor or weak density were removed. The final model consists of *PLCβ3*: 13-92, 99-470, 575-873, with the sidechains of R82, K85, E88, E100, K129, K173, E191, E207, E211, K236, K238, K268, E303, M383, D417, K420, K601, D721, E779, D850, Q853, Y855, and R872 removed due to poor density, *Gα*_*q*_: 38-352 with the sidechains of E49, D69, E70, R73, K98, E104, E115, D117, E125, D155, D165, D169, Q197, Q265, D296, D301, E307, and D321 truncated due to poor density. *Gβ* 1: 6-126 and 133-340, with sidechains from L7, R8, Q9, K15, R19, D20, R22, K23, N36, R46, R52, D66, K89, R96, R134, R137, R214, E215, E226, K280, and R283 truncated due to poor density, *Gγ* 1: 9-61, with the side chains of K20, M21, K29, E47, and E58 truncated due to poor density, *Gβ* 2: 4-126, and 133-340 with sidechains from L7, Q9, R19, K23, E27, Q32, N35, N36, R42, R46, D66, R68, N88, R96, R134, R137, D154, E172, Q175, R214, E215, M216, R251, D258, Q259, K280, K301, D322, and K337 truncated due to poor density, and *Gγ* 2: 8-51 with sidechains from Q11, R13, K14, M21, N24, K29, and K46 truncated due to poor density. Model quality was assessed with validation in Phenix using MolProbity score (49) and geometry evaluation (Table S1). Figures were made using ChimeraX (50, 51).

## Acknowledgements

We thank Chen Zhao for developing and characterizing the ALFA nanobody mediated tethering of G*βγ* to GIRK and for insightful discussions. We thank Venkata S. Mandala for assistance with protein reconstitution and NMR experiments. We thank Christoph A. Haselwandter for comments on the paper. We thank Yi Chun Hsiung for assistance with tissue culture. We thank members of the MacKinnon lab, Jue Chen and members of her lab for helpful discussions. This work was supported by National Institute of General Medical Sciences (NIHF32GM142137 to MEF). RM is an investigator in the Howard Hughes Medical Institute.

We thank Rui Yan and Zhiheng Yu at the HHMI Janelia Cryo-EM Facility for help in microscope operation and data collection. We thank Mark Ebrahim, Johanna Sotiris, and Honkit Ng at the Evelyn Gruss Lipper Cryo-EM Resource Center of Rockefeller University for assistance with cryo-EM screening.

## Data, Materials, and Software Availability

Cryo-EM maps and atomic models for all structures described in this work have been deposited to the Electron Microscopy Data Bank (EMDB) and the Protein Data Bank (PDB), respectively.

## Supporting Information for

**Figure S1:**
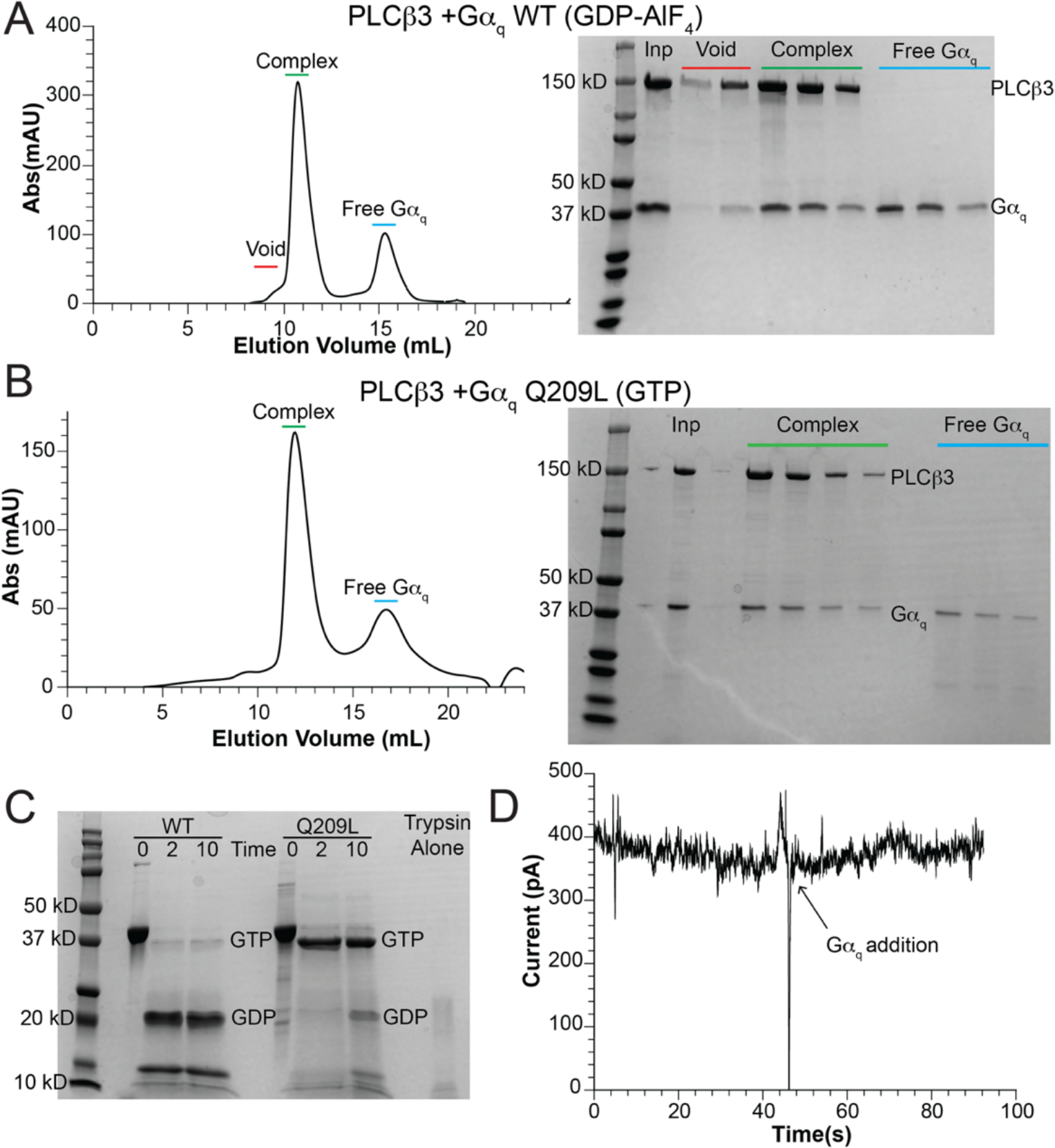
Comparison of *Gα*_*q*_ wildtype and hydrolysis deficient mutant, Q209L. A-B: *PLCβ3-Gα*_*q*_ complex formation with wildtype *Gα*_*q*_ in the presence of GDP-AlF_4_ (A) or *Gα*_*q*_ Q209L in the presence of GTP (B) with size exclusion chromatography profiles of the complex on a Superdex 200 10/300 increase column are on the left and accompanying SDS-PAGE gels are on right. In both cases, *PLCβ3* was mixed with a two-fold molar excess of *Gα*_*q*_, and *Gα*_*q*_ comigrates with *PLCβ3*. Inp refers to the input sample prior to size exclusion chromatography. C: SDS-PAGE gel of trypsin cleavage assay (39) (see methods) to evaluate the nucleotide state of purified *Gα*_*q*_ wildtype (left) or Q209L (right). The time is in minutes and the 0 timepoint is prior to the addition of trypsin. Cleavage of GDP-bound *Gα*_*q*_ produces a band ~ 20 kD and cleavage of GTP-bound *Gα*_*q*_ produces a band ~37 kD (39). Purification of *Gα*_*q*_ Q209L in the presence of GTP yields predominantly GTP-bound protein. D. GIRK current over time before and after addition of 50 *nM Gα*_*q*_ Q209L GTP showing the current is not affected.

**Figure S2:**
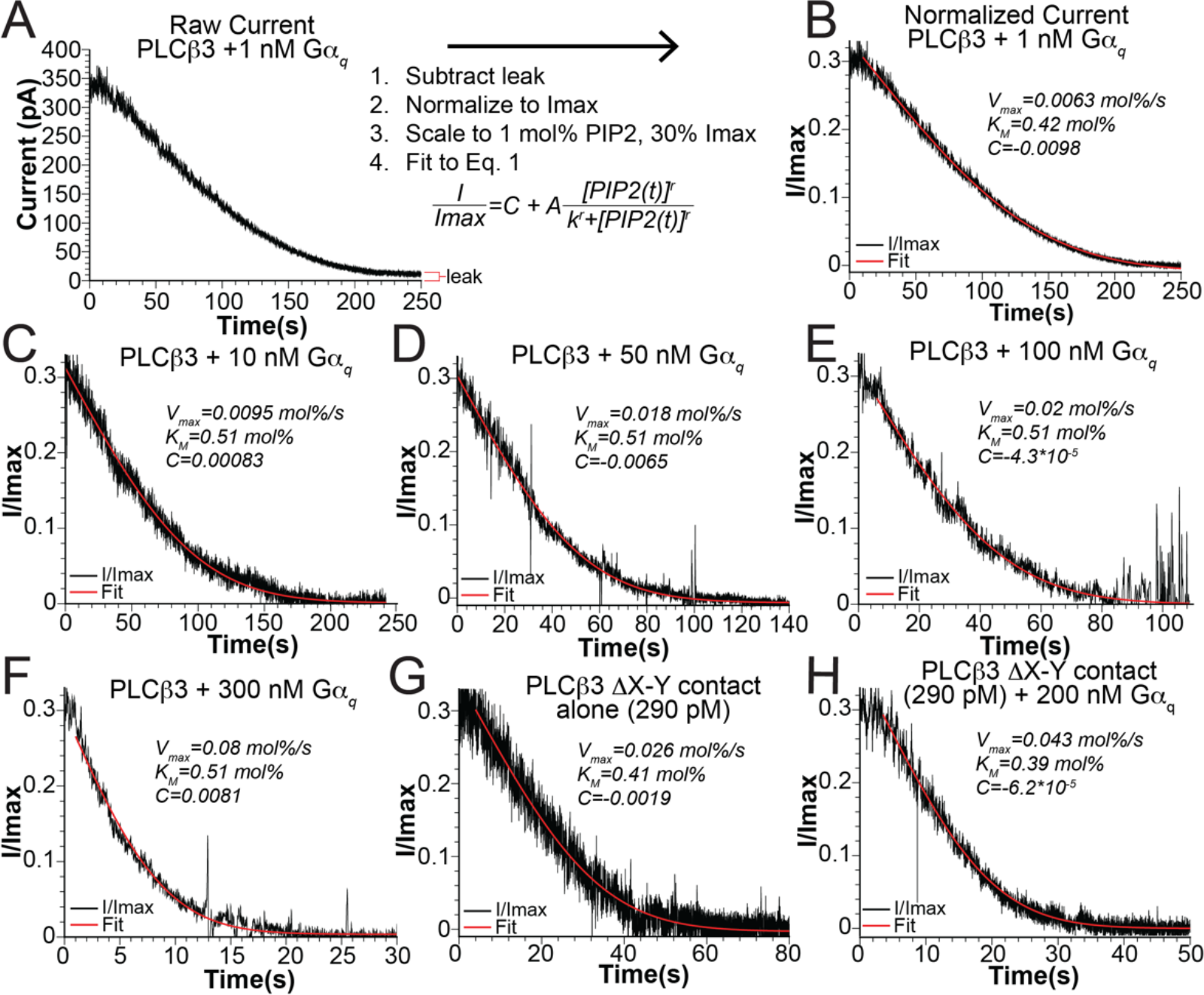
*Gα*_*q*_-dependent activation of *PLCα3*. A: Raw *PLCα3-Gα*_*q*_-induced current decay and analysis scheme. The example shown is in the presence of 1.0 *nM Gα*_*q*_. The red bars in the bottom right corner of the trace demarcate the background leak current that remains in the bilayer when the current decay is complete. This is the background leak current that is subtracted prior to normalization and further analysis. To determine values for *V*_*max*_ and *K*_*M*_, raw currents were leak subtracted, normalized to the maximum current before the decay, *I*_*max*_, and scaled to the starting level of *PIP2* in the bilayer, 1 *mol%*, which is 30% of the maximal GIRK-ALFA current with saturating *PIP2*. The normalized decays were fit to Eq. 8 with free parameters *V*_*max*_, *K*_*M*_, and *C*. B-F: Representative normalized current decay (using 29 *nM* of wildtype *PLCα3*) fit to Eq. 8 to determine *V*_*max*_ and *K*_*M*_ (red curves) in the presence of varying concentrations of *Gα*_*q*_, 1.0 *nM* (B), 10 *nM* (C), 50 *nM* (D), 100 *nM* (E), 300 *nM* (F). H-I: Representative normalized current for *PLCα3* with the structured part of the X-Y linker deleted (*Δ*X-Y contact) using 290 *pM* enzyme fit to Eq. 8 to determine *V*_*max*_ and *K*_*M*_ (red curves) in the absence (H) and presence (I) of 200 *nM Gα*_*q*_.

**Figure S3:**
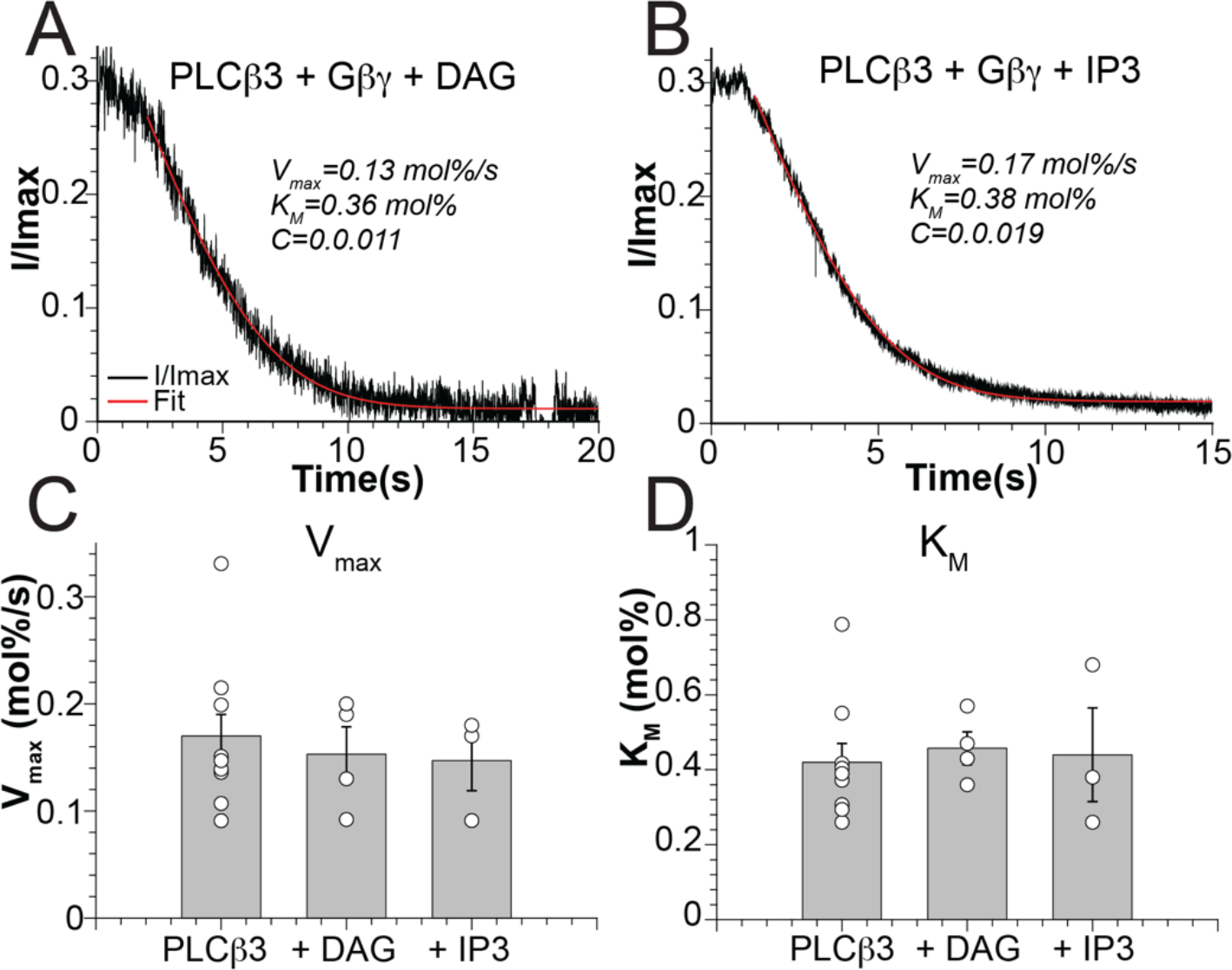
*PLCα3* is not inhibited by the products of its reaction, IP3 and DAG, in our experimental setup. A-B: normalized *PLCα3*-dependent current decays in the presence of *Gαγ* and 1.0 *mol%* DAG (A) or 1.0 *µM* IP3 (B). Red curves are fits to Eq. 8 to determine *V*_*max*_ and *K*_*M*_. R^2^=0.97 (A) and 1.0 (B). C-D: comparison of *V*_*max*_ (C) or *K*_*M*_ (D) in the presence and absence of 1.0 *mol%* DAG and 1.0 *µM* IP3. The bars are the mean, the error bars are standard error of mean, and the circles are values from individual experiments. Data for *PLCα3* alone were reported in (14).

**Figure S4:**
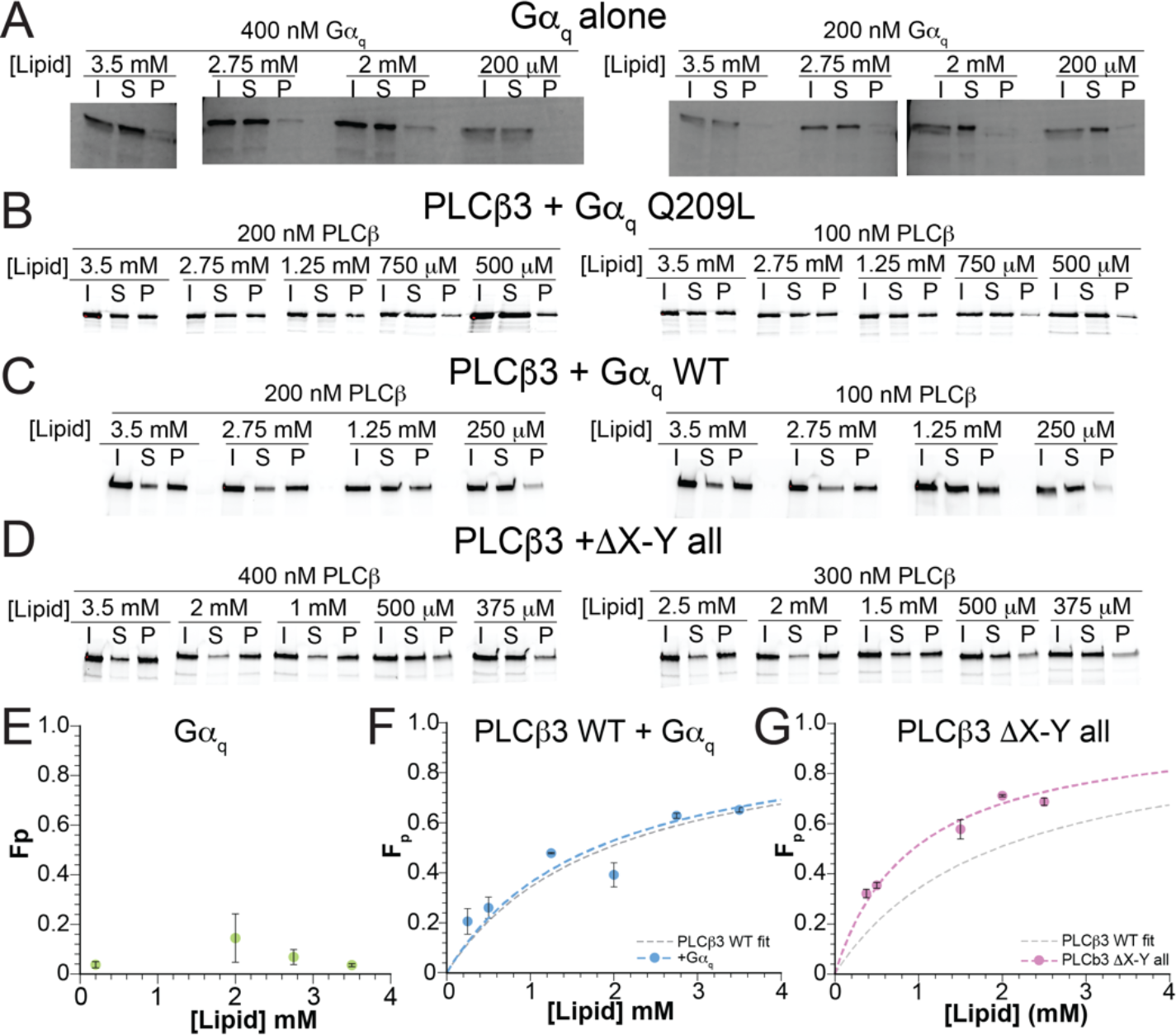
*PLCα3* partitioning in the presence of *Gα*_*q*_. A: Silver stained SDS-PAGE gels of partitioning experiments for wildtype *Gα*_*q*_ with GDP-AlF_4_ (plotted in E). I is input, S is supernatant, and P is pellet. No membrane partitioning behavior is evident as protein in the pellet (P) does not increase with increasing lipid concentration. B-D: SDS-PAGE gels imaged for LD655 fluorescence of wildtype *PLCα3* partitioning experiments in the presence of *Gα*_*q*_ Q209L with GTP (B), wildtype *Gα*_*q*_ GDP-AlF_4_ (C) or *PLCα3 α*X-Y all (D). E-G: Membrane partitioning curve for *Gα*_*q*_ alone (E), wildtype *PLCα3* in the presence of wildtype *Gα*_*q*_ with GDP-AlF_4_ (F-blue) or *PLCα3 α*X-Y all (G-pink) for 2DOPE:1POPC:1POPS LUVs with Fraction Partitioned (F_p_) on the Y axis. Points are the average from 2 repeats for each lipid concentration and error bars are range of mean. Points in E and F were fit to Eq. 5 to determine *K*_*x*_ (dashed blue or pink curve). For wildtype *PLCα3* in the presence of wildtype *Gα*_*q*_, *K*_*x*_=3.1*10^4^, R^2^=0.81. For *PLCα3 α*X-Y all, *K*_*x*_=5.8*10^4^, R^2^=0.96. The fit to Eq. 5 for wildtype *PLCα3* alone is shown as a gray dashed curve for reference (14).

**Figure S5:**
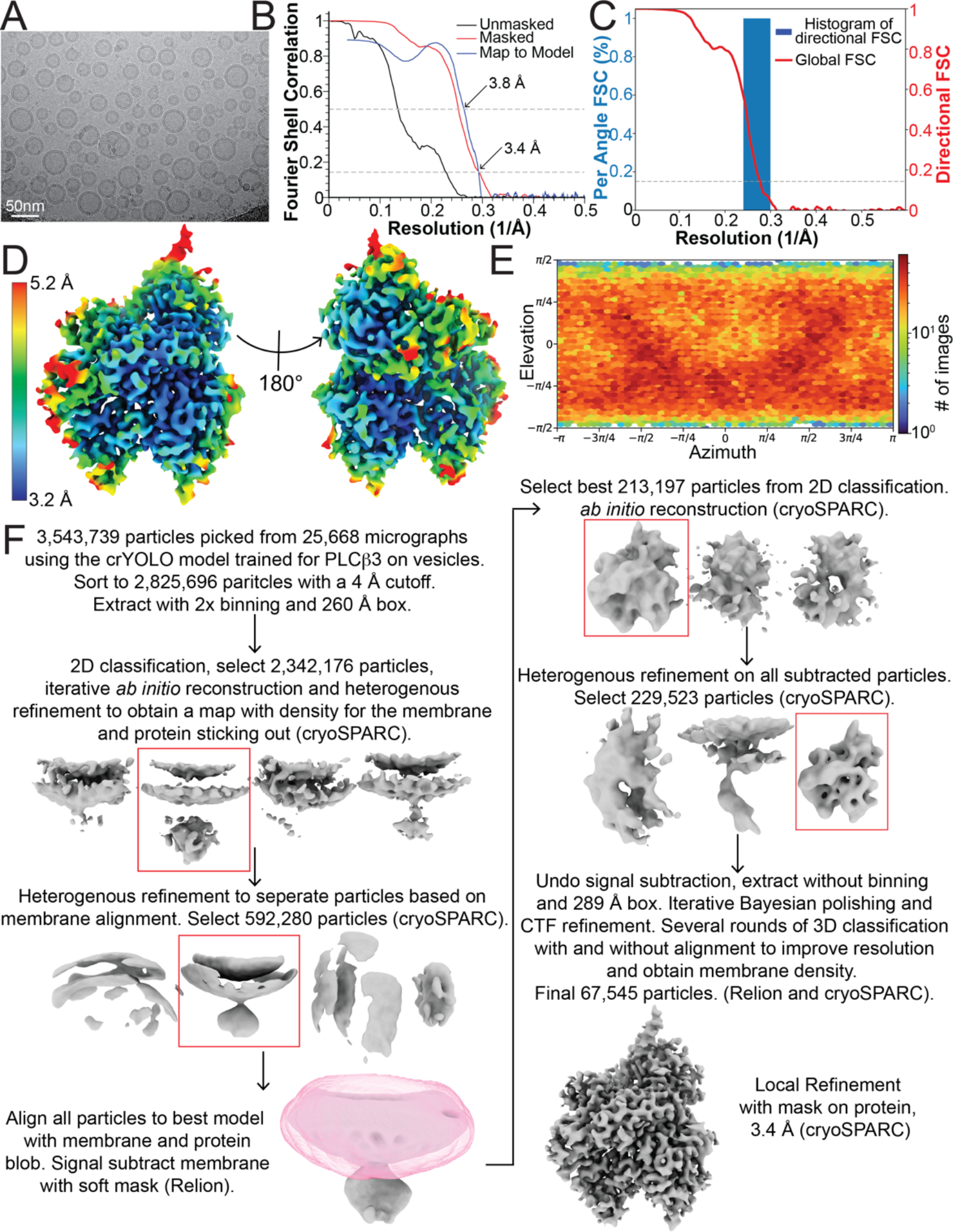
Structure of the *PLCα3-Gα*_*q*_ complex. A: Representative micrograph. B: Fourier shell correlation (FSC) curves for the unmasked (black) and masked (red) maps and between the map and model (blue). The 0.143 and 0.5 thresholds are denoted by dashed lines. C: 3D FSC plot for the final masked map (52). D: Final masked, sharpened map colored by local resolution determined by cryoSPARC. E: Angular distribution plot for the final masked map from cryoSPARC. F: Summary of data processing steps, see methods. Maps shown are unsharpened.

**Figure S6:**
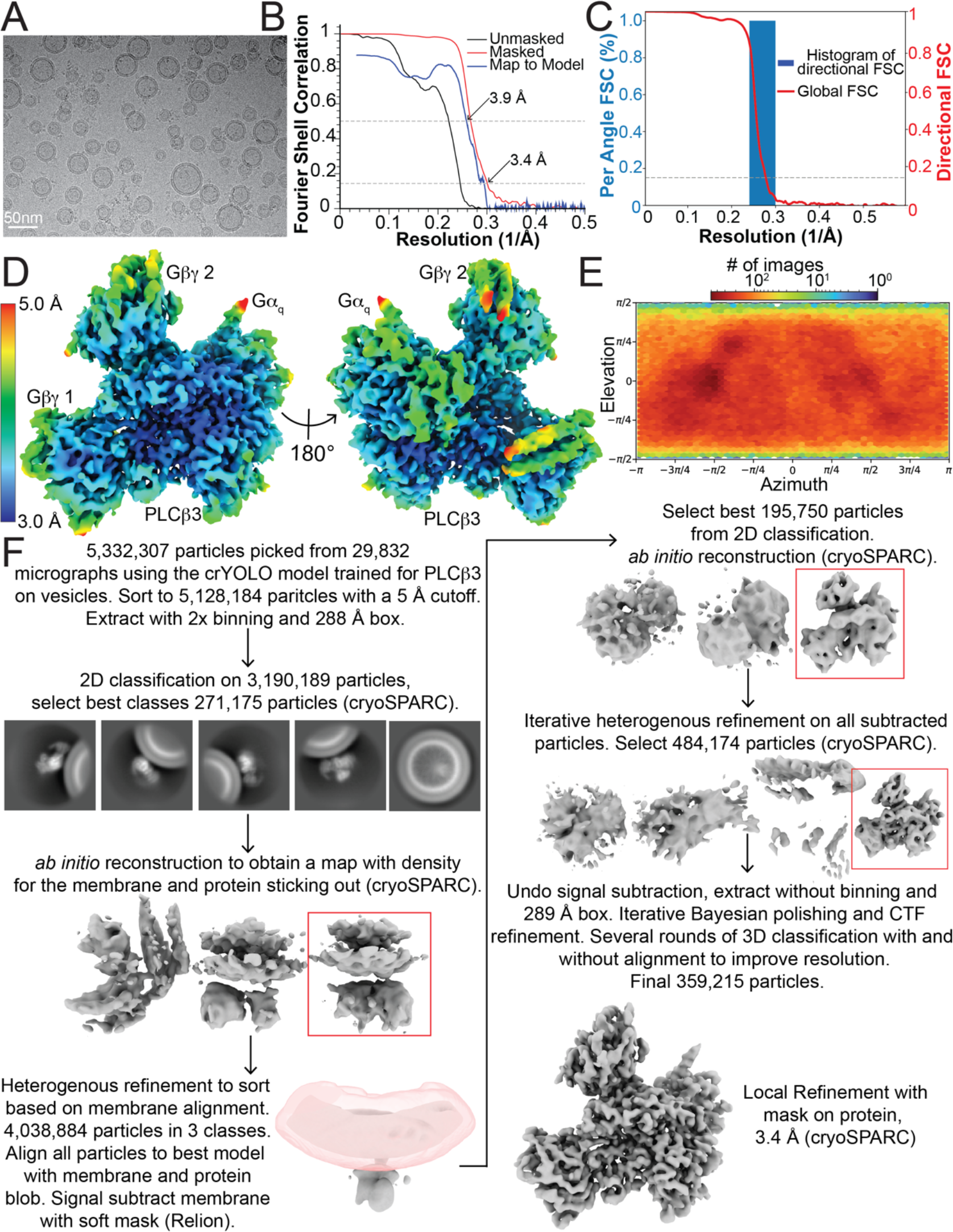
Structure of the *PLCα3-Gαγ(2)-Gα*_*q*_ complex. A: Representative micrograph. B: Fourier shell correlation (FSC) curves for the unmasked (black) and masked (red) maps and between the map and model (blue). The 0.143 and 0.5 thresholds are denoted by dashed lines. C: 3D FSC plot for the final masked map (52). D: Final masked, sharpened map colored by local resolution determined by cryoSPARC. E: Angular distribution plot for the final masked map from cryoSPARC. F: Summary of data processing steps, see methods. Maps shown are unsharpened.

**Figure S7:**
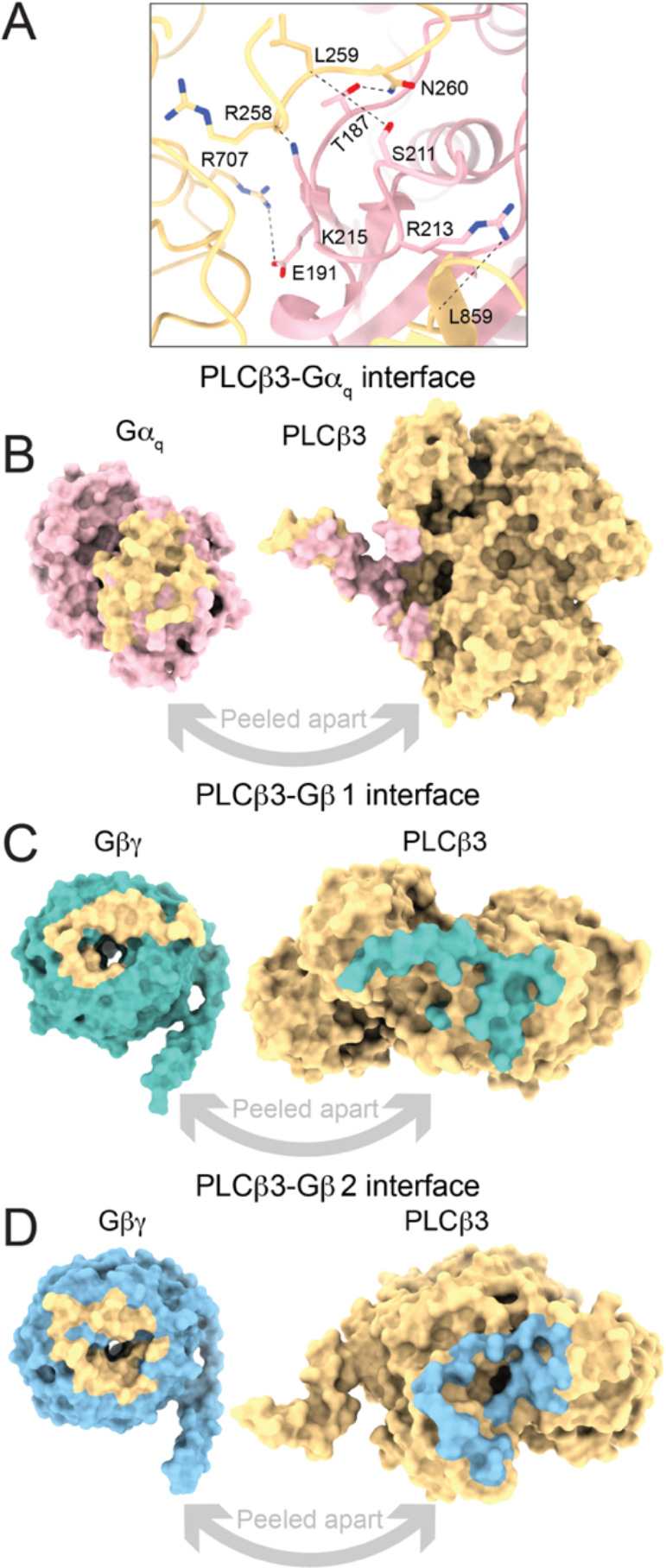
*PLCα3*-G protein interfaces. A: Hydrogen bonds at the *PLCα3-Gα*_*q*_ interface. *PLCα3* is yellow and *Gα*_*q*_ is pink. Relevant side chains are shown as sticks and colored by heteroatom. Hydrogen bonds are denoted by dashed black lines. All hydrogen bonds shown are less than 3.5 Å apart. B-D: Surface representation of the *PLC-Gα*_*q*_ (B), *PLCα3-Gα* 1 (C), or *PLCα3-Gα* 2 (D) interfaces in the *PLCα3-Gαγ(2)-Gα*_*q*_ complex peeled apart to show extensive interactions. *PLCα3* is yellow, *Gα*_*q*_ is pink, *Gα* 1 is dark teal, and *Gα* 2 is light blue. *Gγ* 1 and 2 were omitted for clarity. Residues on *PLCα3* that interact with G proteins are colored according to the corresponding G protein and residues on the G proteins that interact with *PLCα3* are colored in yellow. Interface residues were determined using the ChimeraX interface feature using a buried surface area cutoff of 15 Å^2^. Interfaces are comparable to structures determined with each G protein on its own.

**Figure S8:**
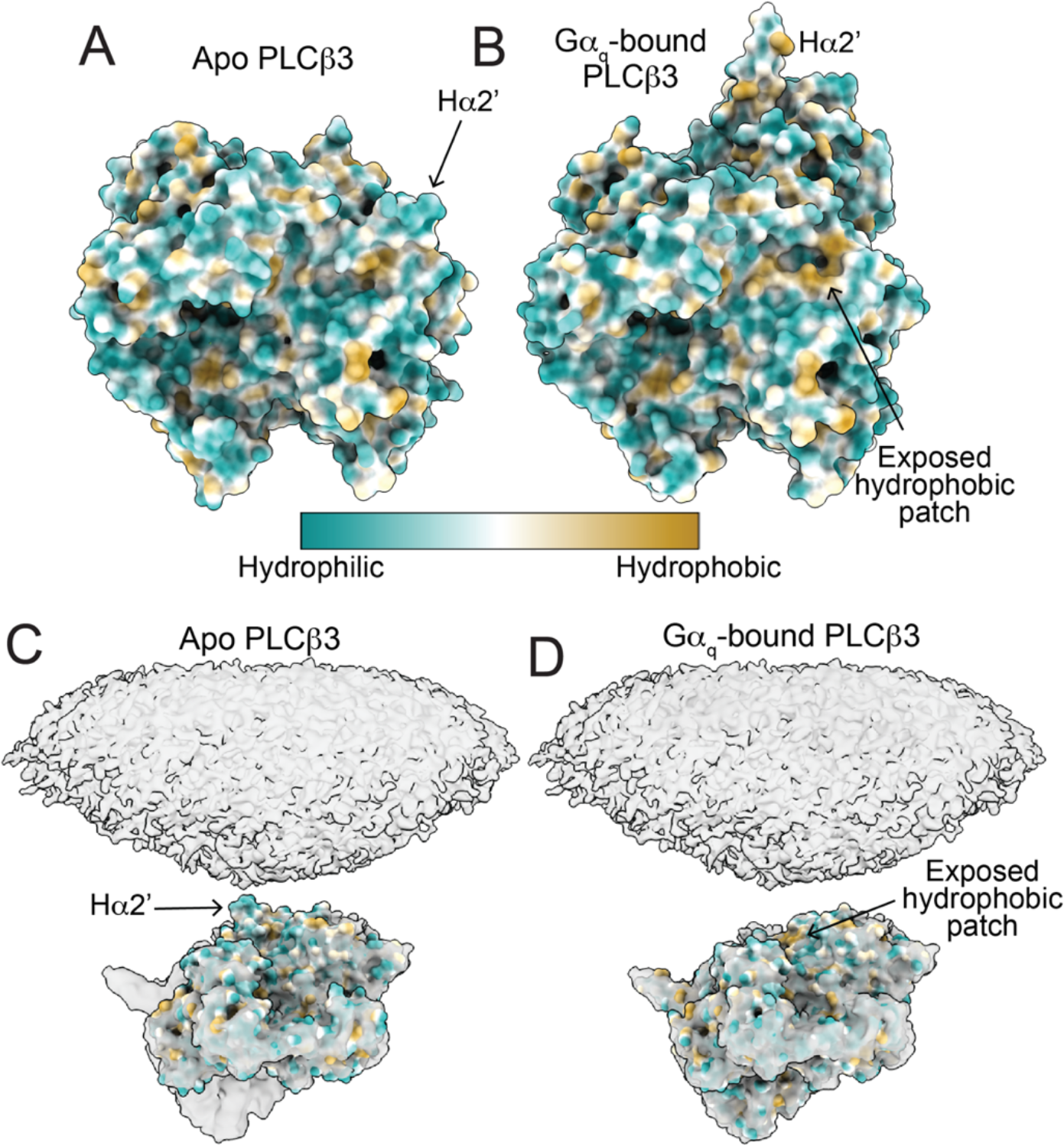
Potential role of the H*α*2’ in membrane association of the *PLCα3* catalytic core. A-B: Catalytic core of apo (A) or *Gα*_*q*_-bound (B) *PLCα3* colored by hydrophobicity highlighting a hydrophobic patch covered by H*α*2’ in the apo conformation. C-D: Hydrophobicity-colored apo (C) or *Gα*_*q*_-bound (D) *PLCα3* catalytic core fit into the density of the membrane associated *PLCα3-Gα*_*q*_ complex. The exposed hydrophobic patch is close to the position of membrane association in the *Gα*_*q*_-bound conformation and the H*α*2’ associated with the catalytic core could hinder membrane association.

**Table S1:** Cryo-EM collection parameters and model statistics.

| Collection Parameters | *PLCβ3-Gα<sub>q</sub>* complex | *PLCβ3-Gβγ(2)-Gα<sub>q</sub>* complex |
| --- | --- | --- |
| Accelerating Voltage (kV) | 300 | 300 |
| Number of frames | 50 | 50 |
| Dose (e<sup>-</sup>/Å<sup>2</sup>) | 60 | 60 |
| Defocus Range (μm) | 1.5-2.5 | 1.5-2.5 |
| Exposure Time (s) | 4.26 | 3.86152 |
| Original Pixel size (Å) | 0.4195 | 0.4195 |
| Map Parameters |  |  |
| Final Pixel Size (Å) | 0.839 | 0.839 |
| Symmetry | C1 | C1 |
| Total micrographs | 25,668 | 29,832 |
| Initial particles | 3,543,739 | 5,332,307 |
| Final particles | 67,545 | 359,215 |
| Map resolution (Å) | 3.43 | 3.37 |
| FSC threshold | 0.143 | 0.143 |
| Map local resolution range (Å) | 3.2-5.2 | 3-5 |
| Map sharpening B factor (Å<sup>2</sup>) | -119.5 | -137 |
| Initial Model Used | 4GNK | 8EMW |
| Model resolution (Å) (FSC<sub>model</sub>=0.5) | 3.8 | 3.9 |
| Model Composition |  |  |
| Nonhydrogen atoms | 8,553 | 13,960 |
| Protein residues | 1069 | 1823 |
| Ligands | 4 | 4 |
| r.m.s. deviations bond length (Å) | 0.008 | 0.011 |
| r.m.s. deviations bond angles (°) | 0.804 | 1.092 |
| Validation |  |  |
| MolProbity Score | 1.71 | 1.99 |
| Clash Score | 6.67 | 9.24 |
| Poor Rotamers (%) | 0 | 0.07 |
| Ramachandran Plot |  |  |
| Favored (%) | 95.0 | 91.68 |
| Allowed (%) | 5.0 | 8.32 |
| Disallowed (%) | 0 | 0 |

**Table S2:** *PLCα3*-G protein interface buried area comparison.

| *Complex* | *Gβγ 1 interface (Å<sup>2</sup>)* | *Gβγ 2 interface (Å<sup>2</sup>)* | *Gα<sub>q</sub> interface (Å<sup>2</sup>)* |
| --- | --- | --- | --- |
| *PLCβ-Gβγ<sup>f</sup>* | 807 | 1105 | - |
| *PLCβ-Gα<sub>q</sub>* | - | - | 1467 |
| *PLCβ-Gβγ(2)-Gα<sub>q</sub>* | 827 | 1001 | 1482 |

1. Previously reported (14)

**Table S3:** *PLCα3*-G protein interface residue comparison. Interface residues were determined using the ChimeraX interface feature using a buried surface area cutoff of 15 Å^2^.

| *Complex* | *Gβγ1 interface (Å<sup>2</sup>)* | *Gβγ2 interface (Å<sup>2</sup>)* | *Gα<sub>q</sub> interface (Å<sup>2</sup>)* |
| --- | --- | --- | --- |
| *PLCβ-Gβγ<sup>I</sup>* | 34 | 44 | - |
| *PLCβ-Gα<sub>q</sub>* | - | - | 56 |
| *PLCβ-Gβγ(2)-Gα<sub>q</sub>* | 35 | 44 | 56 |

1. Previously Reported (14)

## Appendix 1 The integrated Michaelis-Menten rate equation

We did not invent the analysis of enzyme kinetics using the time evolution of product appearance (or substrate disappearance). Indeed, Michaelis and Menten in their original study of invertase, and many others since, analyzed the time course of enzyme activity, but originally with a different mathematical formalism because modern numerical integration methods were not available (53, 54). The Michaelis-Menten rate equation has been popularized and is the approach almost exclusively taught in books and classrooms. For the approach we have taken, we thought it should be sufficient to reference excellent published explanations (53, 54), however, recurrent misunderstanding during the review process of our work on this subject prompted us to demonstrate here with a simple example of how it works. This exercise is not intended to be a further analysis of our data, but a simple demonstration of how, for the system and conditions we are studying, the integrated Michaelis-Menten equation allows us to approximate *K*_*M*_ and *V*_*max*_ (and therefore, with independent knowledge of enzyme concentration, which we have, *k*_*cat*_). We begin with the reaction scheme:

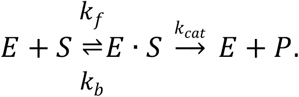

In our case *E* is *PLCα3, S* is *PIP2, E* • *S* stands for a molecular complex composed of one *E* and one *S* molecule, and *P* stands for the products *IP3* and *DAG*. The evolution of this system from *t* = 0 is given by the solution of the equations:

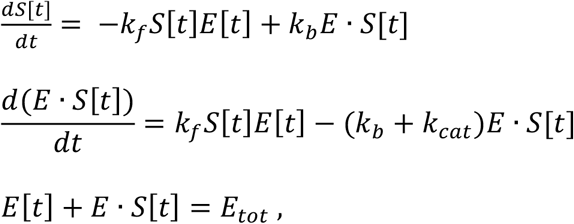

with initial conditions *S*[0] = 1, *E*[0] = *E*_*tot*_. We refer to the solution of these equations for *S[t]* (i.e., the *PIP2* concentration as a function of time) as the ‘true’ solution.

Next, consider the approach we take in the main text of this paper. We express the rate of substrate (*PIP2*) disappearance (requiring a minus sign) using the Michaelis-Menten rate equation,

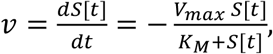

which, for *K*_*M*_ = (*k*_*b*_ + *k*_*cat*_)/*k*_*f*_ and *V*_*max*_ = *k*_*cat*_ *E*_*tot*_, can be calculated for the reaction scheme above if *k*_*f*_*S*[*t*]*E*[*t*] = (*k*_*b*_ + *k*_*cat*_)*E* • *S*[*t*]. We then integrate *v* from time 0 to *t* to find

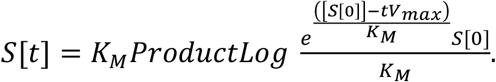

We refer to the above for *S[t]* as the ‘approximate’ solution. Next, we assign rate constant values, calculate the ‘true’ solution (i.e., by solving the above rate equations) and then we fit *S[t]* to the ‘true’ solution to get an ‘approximate’ solution. For *k*_*f*_ = 10,000 (*mol% s*)^−1^, *k*_*b*_ = 4000 *s*^*-1*^, with *k*_*cat*_ = 1.7 *s*^*-1*^ and *E*_*tot*_ = 0.0015 *mol%* (approximating wild type, panel A), with *k*_*cat*_ = 2000 *s*^*-1*^ and *E*_*tot*_ = 0.000015 *mol%* (approximating an X-Y mutant *PLCα3*, panel B) or with *k*_*cat*_ = 6.6 *s*^*-1*^ and *E*_*tot*_ = 0.045 (approximating saturating *Gαγ* + 10 *nM Gα*_*q*_, *panel C*) the results are as follows (black curves denote ‘true’ solutions and red dashed curves denote the fitted, approximate solutions):

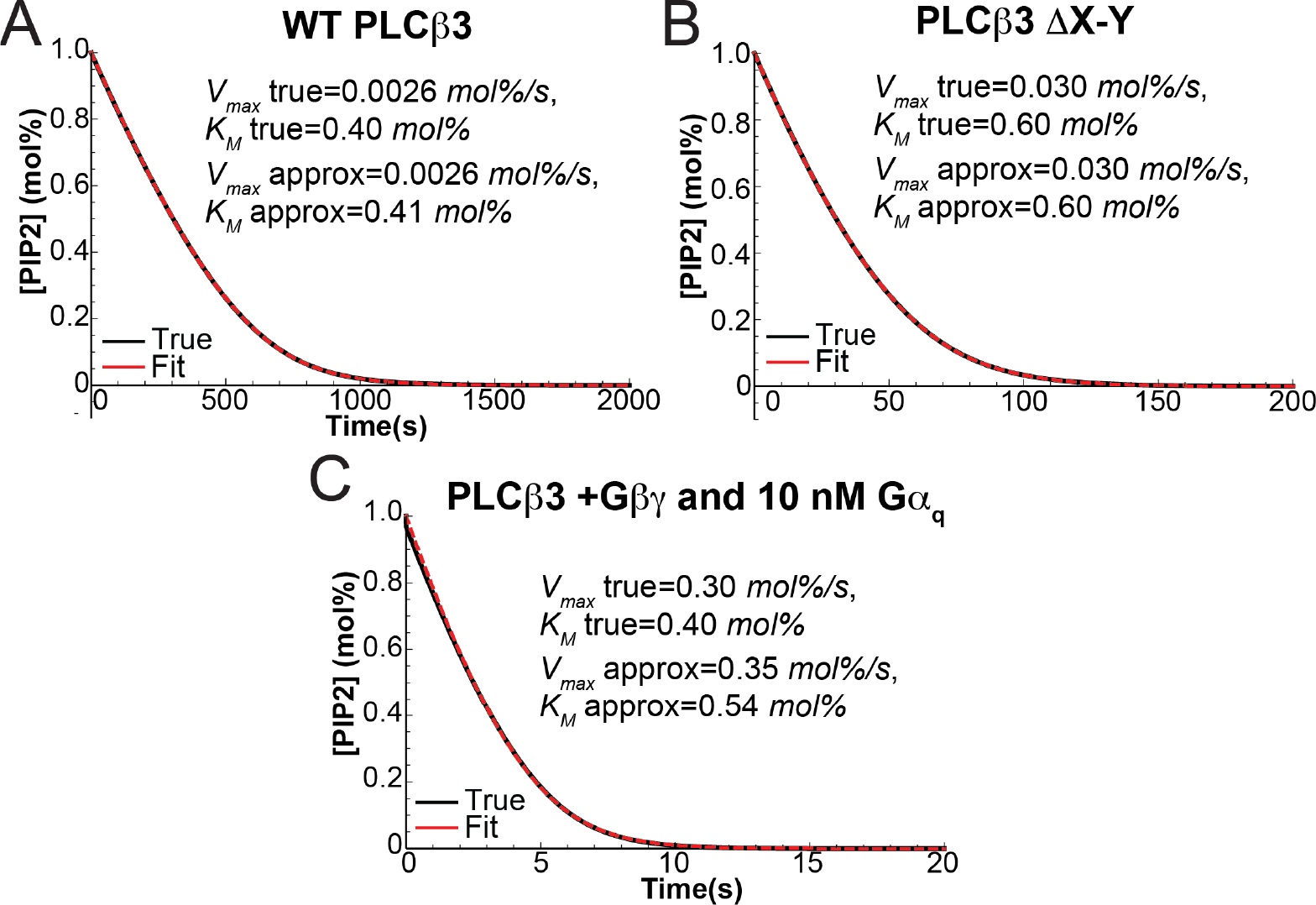

The integrated form of the Michaelis-Menten equation (Lambert W Function) approximates the ‘true’ process (as described by the reaction equations above that define the system) for a wide range of *k*_*f*_, *k*_*b*_ and *k*_*cat*_ values. As when using the Michaelis-Menten initial rate equation, it is important that *E*_*tot*_ ≪*S[0]* + *K*_*M*_, which is the case for our studies. For the conditions shown in Panel C, there is a deviation between the fitted and true values of *K*_*M*_ and *V*_*max*_. These conditions correspond to our experiments in which a saturating quantity of *Gαγ* is used, which increases the local enzyme concentration. Even in this most extreme case, *E*_*tot*_ ≪*S[0]* + *K*_*M*_ remains true, and the expected inaccuracy is small, likely within the uncertainty of our experimental data.

The reader should realize that the only real difference between measuring initial rates at various substrate concentrations and measuring the slope along the decay curve (which is then paired with the substrate concentration on the Y-axis) is the presence of product in the latter and the absence of product in the former. These approaches should give the same result if the products do not alter the function of the enzyme. For *PLCα3* we do not observe product inhibition (Fig. S3).

## Appendix 2 A calculation to explore whether the high *k*_*cat*_ in the X-Y linker deletions renders *PLCα3* insensitive to *Gα*_*q*_ regulation

In this study we demonstrated that *Gα*_*q*_ increases *k*_*cat*_ by about 35-fold. We further demonstrated that when the X-Y linker is removed by mutation, *Gα*_*q*_ no longer causes the 35-fold enhancement. From these observations, we hypothesized that the effect of *Gα*_*q*_ to increase *k*_*cat*_ is somehow mediated by the X-Y linker. Because X-Y linker removal increases *k*_*cat*_ from 1.7 *s*^*-1*^ to ~ 2000 *s*^*-1*^, here we ask the question, could the mutant *PLCα3* have entered a diffusion-limited realm? In this case enzyme turnover would become insensitive to *k*_*cat*_ enhancement owing to a shift in the ‘rate limiting step’ to diffusion of substrate up to the active site, and our hypothesis that *Gα*_*q*_ regulates the active site would be incorrect. To assess the likelihood of this possibility, we present the following calculation.

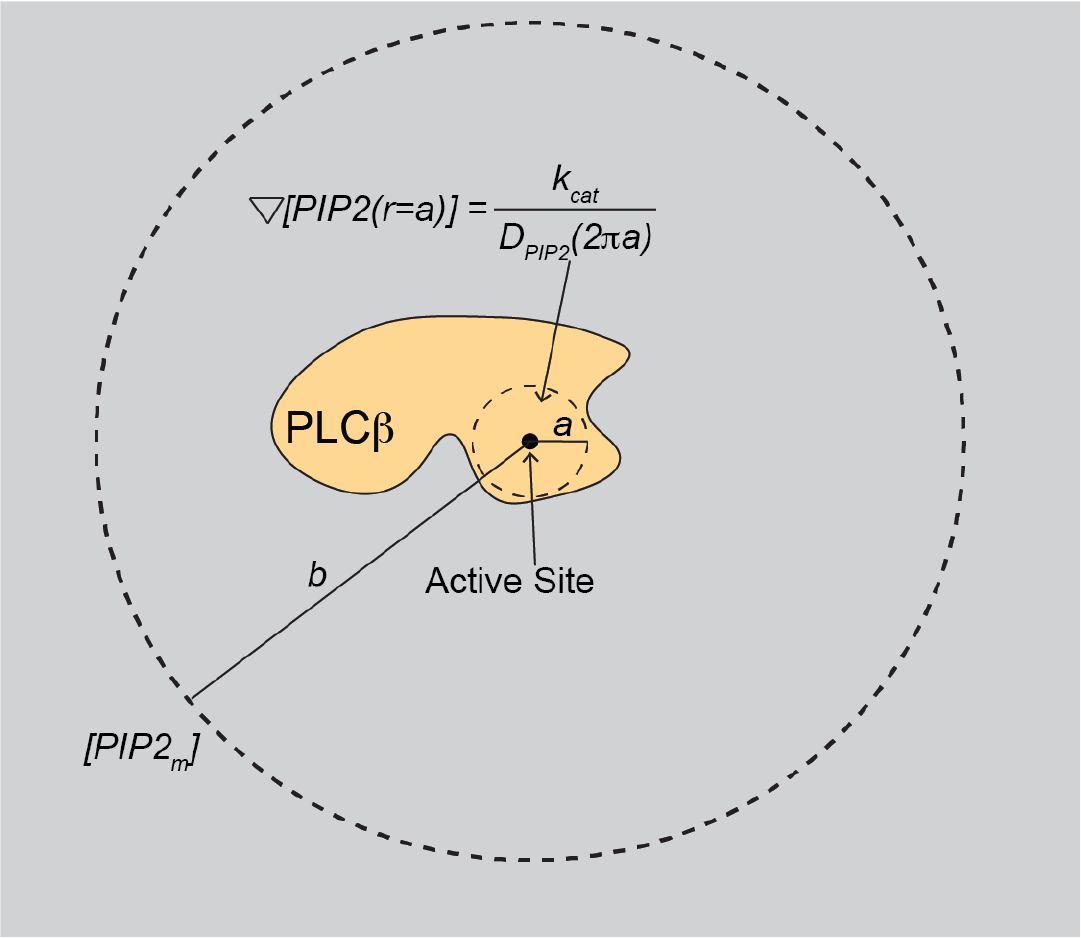

Imagine a *PLCα3* enzyme on the membrane surface and define an imaginary circle around its active site with a radius *a* = 1 *nm* (shown above). Given the concentration of *PIP2* in the membrane and its diffusion coefficient *D*_*PIP2*_ (~ 1 *µm*^*2*^ *s*^*-1*^) we can calculate the concentration profile of *PIP2* as a function of radial distance *r* away from the active site when *k*_*cat*_ equals its value in the absence of the X-Y linker, say 2000 *s*_*-1*_, to assess whether *PIP2* becomes depleted near the active site, i.e., diffusion limited. An important assumption must be made in this calculation concerning boundary conditions on steady state diffusion in 2 dimensions. We note that *PLCα3* also diffuses (say, with *D*_*PLCβ3*_ = 1 *µm*^*2*^ *s*^*-1*^), and therefore keeps moving a distance approximately 63 *nm* every 1 *ms*. Our assumption is, *PLCα3* ‘refreshes’ its concentration at boundary *b*, 63 *nm* away. Thus, we will create an imagined steady state *PIP2* concentration profile as a function of distance *r* away from the active site, between ‘bulk’ membrane concentration at *b* and a gradient of concentration at *a* that will yield a net inward flux of *PLCα3* equal to *k*_*cat*_. The concentration profile is given by solving

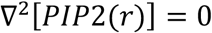

subject to the boundary conditions

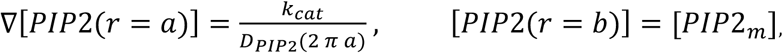

which gives

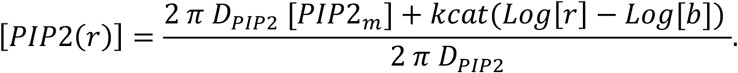

Solutions to this expression are shown in the two graphs below.

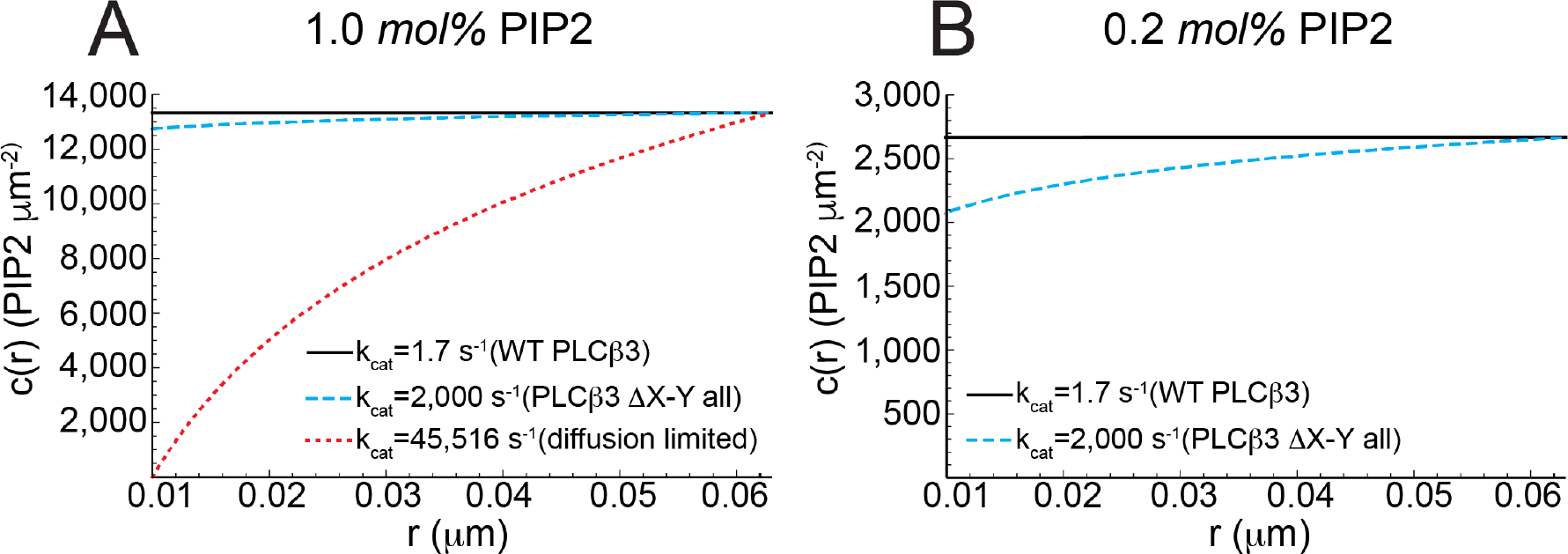

Panel A shows, with 1.0 *mol% [PIP2*_*m*_] (~13,333 *PIP2 µm*^*-2*^), the calculated concentration profile for *k*_*cat*_ = 1.7 *s*^*-1*^ (black), 2000 *s*^*-1*^ (blue) and 45,516 *s*^*-1*^ (red, the diffusion limited case, corresponding to the boundary condition [*PIP*2(*r* = *a*)] = 0) with *D*_*PIP2*_ = 1.0 *µm*^*2*^ *s*^*-1*^. At 2000 *s*^*-1*^, minimal depletion of substrate occurs. Panel B shows, with 0.2 *mol% [PIP2*_*m*_], the concentration profile for *k*_*cat*_ = 1.7 *s*^*-1*^ (black) and 2000 *s*_*-1*_ (blue). Even at this lower *PIP2* concentration, the reaction is far from diffusion limited. While several assumptions are made in this calculation, it presents a rational argument that the *PLCα3* reaction under our assay conditions are far from diffusion limited. While a rate increase from 2000 *s*_*-1*_ to 35 times 2000 *s*^*-1*^ (The fold activation by *Gα*_*q*_) is not possible based on the diffusion limited example above, we should see a substantial rate increase, but we do not. For this reason, we hypothesize that the increased *k*_*cat*_, from 1.7 *s*^*-1*^ to ~60 *s*^*-1*^, due to binding of *Gα*_*q*_ to *PLCα3*, results from allosteric regulation mediated at least in part through the X-Y linker.

## References

1. R. Rodnight, Cerebral diphosphoinositide breakdown: activation, complexity and distribution in animal (mainly nervous) tissues. Biochemical Journal 63, 223–231 (1956).

2. P. Kemp, G. Hübscher, J. Hawthorne, Phosphoinositides. 3. Enzymic hydrolysis of inositol-containing phospholipids. Biochemical Journal 79, 193–200 (1961).

3. C. V. Robinson, T. Rohacs, S. B. Hansen, Tools for Understanding Nanoscale Lipid Regulation of Ion Channels. Trends in Biochemical Sciences 44, 795–806 (2019).

4. J. A. Poveda et al., Lipid modulation of ion channels through specific binding sites. Biochimica et Biophysica Acta - Biomembranes 1838, 1560–1567 (2014).

5. S. B. Hansen (2015) Lipid agonism: The PIP2 paradigm of ligand-gated ion channels. in Biochimica et Biophysica Acta - Molecular and Cell Biology of Lipids (Elsevier B.V.), pp 620–628.

6. G. Kadamur, E. M. Ross (2013) Mammalian Phospholipase C. in Annual Review of Physiology, pp 127–154.

7. M. J. Berridge, Inositol Trisphosphate and Diacylglycerol: Two Interacting Second Messengers. Annual Review of Biochemistry 56, 159–193 (1987).

8. A. M. Lyon, J. J. G. Tesmer (2013) Structural insights into phospholipase C-β function. in Molecular Pharmacology, pp 488-500.

9. A. M. Lyon, J. A. Begley, T. D. Manett, J. J. G. Tesmer (2014) Molecular mechanisms of phospholipase C β3 autoinhibition. in Structure, pp 1844–1854.

10. A. M. Lyon, S. Dutta, C. A. Boguth, G. Skiniotis, J. J. G. Tesmer (2013) Fulllength Gαq-phospholipase C-β3 structure reveals interfaces of the C-terminal coiled-coil domain. in Nature Structural and Molecular Biology (Nature Publishing Group), pp 355–362.

11. A. M. Lyon et al. (2011) An autoinhibitory helix in the C-terminal region of phospholipase C-β mediates Gαq activation. in Nature Structural and Molecular Biology (Nature Publishing Group), pp 999–1005.

12. S. N. Hicks et al. (2008) General and Versatile Autoinhibition of PLC Isozymes. in Molecular Cell, pp 383–394.

13. T. H. Charpentier et al. (2014) Membrane-induced allosteric control of phospholipase C-β isozymes. in Journal of Biological Chemistry, pp 29545–29557.

14. Maria E. Falzone, R. MacKinnon, <i>Gβγ</i> activates <i>PIP2</i> hydrolysis by recruiting and orienting <i>PLCβ</i> on the membrane surface. Proceedings of the National Academy of Sciences 120, e2301121120 (2023).

15. J. R. Hepler et al. (1996) Functional importance of the amino terminus of Gqα. In Journal of Biological Chemistry, pp 496–504.

16. G. L. Waldo et al. (2010) Kinetic Scaffolding Mediated by a Phospholipase C–b and Gq Signaling Complex. in Science.

17. R. A. Rebres et al. (2011) Synergistic Ca2+ responses by Gαi- and Gαq-coupled G-protein-coupled receptors require a single PLCβ isoform that is sensitive to both Gβγ and Gαq. in Journal of Biological Chemistry, pp 942–951.

18. T. I. A. Roach et al. (2008) Signaling and cross-talk by C5a and UDP in macrophages selectively use PLCβ3 to regulate intracellular free calcium. in Journal of Biological Chemistry, pp 17351–17361.

19. F. Philip, G. Kadamur, R. G. Silos, J. Woodson, E. M. Ross (2010) Synergistic activation of phospholipase C-β3 by Gαq and Gβγ describes a simple two-state coincidence detector. in Current Biology, pp 1327–1335.

20. M. Goličnik, On the Lambert W function and its utility in biochemical kinetics. Biochemical Engineering Journal 63, 116–123 (2012).

21. M. J. W. Adjobo-Hermans et al. (2013) PLCβ isoforms differ in their subcellular location and their CT-domain dependent interaction with Gαq. in Cellular Signalling (Elsevier Inc.), pp 255–263.

22. D. E. Bosch et al., A P-loop Mutation in Gα Subunits Prevents Transition to the Active State: Implications for G-protein Signaling in Fungal Pathogenesis. PLOS Pathogens 8, e1002553 (2012).

23. W. Huang et al. (2018) A membrane-associated, fluorogenic reporter for mammalian phospholipase C isozymes. in Journal of Biological Chemistry, pp 1728–1735.

24. M. Maziarz et al., Atypical activation of the G protein Gαq by the oncogenic mutation Q209P. Journal of Biological Chemistry 293, 19586–19599 (2018).

25. S. H. White, W. C. Wimley, A. S. Ladokhin, K. Hristova, “[4] Protein folding in membranes: Determining energetics of peptide-bilayer interactions” in Methods in Enzymology. (Academic Press, 1998), vol. 295, pp. 62–87.

26. A. M. Lyon, V. G. Taylor, J. J. G. Tesmer (2014) Strike a pose: Gαq complexes at the membrane. in Trends in Pharmacological Sciences (Elsevier Ltd), pp 23–30.

27. J. Won et al., Molecular architecture of the Gαi-bound TRPC5 ion channel | Nature Communications. Nature Communications 14, 2550 (2023).

28. C. Qi et al., Structural basis of adenylyl cyclase 9 activation. Nature Communications 13, 1045 (2022).

29. B. Khanppnavar et al. (2023) Structural basis of activation and inhibition of the Ca2+/calmodulin-sensitive adenylyl cyclase 8. (bioRxiv).

30. C. Qi, S. Sorrentino, O. Medalia, V. M. Korkhov, The structure of a membrane adenylyl cyclase bound to an activated stimulatory G protein. Science 364, 389–394 (2019).

31. J. Ma, L. Weng, B. C. Bastian, X. Chen, Functional characterization of uveal melanoma oncogenes. Oncogene 40, 806–820 (2021).

32. H. T. N. Phan, N. H. Kim, W. Wei, G. G. Tall, A. V. Smrcka, Uveal melanoma– associated mutations in PLCβ4 are constitutively activating and promote melanocyte proliferation and tumorigenesis. Science Signaling 14, eabj4243 (2021).

33. J. J. Park et al., Oncogenic signaling in uveal melanoma. Pigment Cell & Melanoma Research 31, 661–672 (2018).

34. E. E. Garland-Kuntz et al. (2018) Direct observation of conformational dynamics of the PH domain in phospholipases Cϵ and β may contribute to subfamily-specific roles in regulation. in Journal of Biological Chemistry, pp 17477–17490.

35. K. Muralidharan, M. M. Van Camp, A. M. Lyon, Structure and regulation of phospholipase Cβ and ε at the membrane. Chemistry and Physics of Lipids 235, 105050 (2021).

36. Q. Zheng et al., Electronic tuning of self-healing fluorophores for live-cell and single-molecule imaging. Chem. Sci. 8, 755–762 (2016).

37. C. Zhao, R. MacKinnon, Structural and functional analyses of a GPCR-inhibited ion channel TRPM3. Neuron 0 (2022).

38. P. Y. Chan et al., Purification of heterotrimeric G protein α subunits by GST-Ric-8 association: Primary characterization of purified Gαolf. Journal of Biological Chemistry 286, 2625–2635 (2011).

39. E. P. Marin et al., The Function of Interdomain Interactions in Controlling Nucleotide Exchange Rates in Transducin. Journal of Biological Chemistry 276, 23873–23880 (2001).

40. P. Chidiac, V. S. Markin, E. M. Ross, Kinetic control of guanine nucleotide binding to soluble Gα(q). Biochemical Pharmacology 58, 39–48 (1999).

41. D. N. Mastronarde, Automated electron microscope tomography using robust prediction of specimen movements. Journal of Structural Biology 152, 36–51 (2005).

42. J. Zivanov et al., New tools for automated high-resolution cryo-EM structure determination in RELION-3. eLife 7, e42166 (2018).

43. A. Rohou, N. Grigorieff, CTFFIND4: Fast and accurate defocus estimation from electron micrographs. Journal of Structural Biology 192, 216–221 (2015).

44. S. Q. Zheng et al., MotionCor2: Anisotropic correction of beam-induced motion for improved cryo-electron microscopy. Nature Methods 14, 331–332 (2017).

45. T. Wagner et al., SPHIRE-crYOLO is a fast and accurate fully automated particle picker for cryo-EM. Communications Biology 2, 1–13 (2019).

46. J. Zivanov, T. Nakane, S. H. W. Scheres, A Bayesian approach to beam-induced motion correction in cryo-EM single-particle analysis. IUCrJ 6, 5–17 (2019).

47. J. Zivanov et al., RELION-3 : new tools for automated high-resolution cryo-EM structure determination. bioRxiv 10.1101/421123, 1-38 (2018).

48. P. V. Afonine, J. J. Headd, T. C. Terwilliger, P. D. Adams, PHENIX News. Computational Crystallography Newsletter 4, 43–44 (2013).

49. V. B. Chen et al., MolProbity: All-atom structure validation for macromolecular crystallography. Acta Crystallographica Section D: Biological Crystallography 66, 12–21 (2010).

50. E. F. Pettersen et al., UCSF Chimera - A visualization system for exploratory research and analysis. Journal of Computational Chemistry 25, 1605–1612 (2004).

51. E. F. Pettersen et al., UCSF ChimeraX: Structure visualization for researchers, educators, and developers. Protein Science 30, 70–82 (2021).

52. Y. Z. Tan et al., Addressing preferred specimen orientation in single-particle cryo-EM through tilting. Nature Methods 14, 793–796 (2017).

53. M. Goličnik, The integrated Michaelis-Menten rate equation: <i>déjà vu</i> or <i>vu jàdé</i> ? Journal of Enzyme Inhibition and Medicinal Chemistry 28, 879–893 (2013).

54. K. A. Johnson, R. S. Goody, The Original Michaelis Constant: Translation of the 1913 Michaelis–Menten Paper. Biochemistry 50, 8264–8269 (2011).

